# Engineering a cytochrome P450 *O*-demethylase for the bioconversion of hardwood lignin

**DOI:** 10.64898/2026.02.18.706646

**Authors:** Megan E. Wolf, Daniel J. Hinchen, Michael Zahn, John E. McGeehan, Lindsay D. Eltis

## Abstract

Lignin is a promising alternative to petroleum as a feedstock for the chemical industry. Emergent strategies for lignin valorization involve tandem processes in which biomass is chemocatalytically fractionated followed by biotransformation of the depolymerized lignin by microbial cell factories. A rate-limiting step in this biotransformation is *O*-demethylation of the lignin-derived monomers. The reductive catalytic fractionation of hardwood biomass generates high yields of two classes of monomers: 4-alkylguaiacols and 4-alkylsyringols. To better understand the biotransformation of these monomers, we studied AgcA, a cytochrome P450, and AgcB, the cognate reductase, that together catalyze the *O*-demethylation of 4-alkylguaiacols. A 1.82 Å resolution crystal structure of AgcA_EP4_ from *Rhodococcus rhodochrous* EP4 in complex with 4-ethylguaiacol identified residues Leu78, Ala293 and Phe166 as potential specificity determinants. Substitution of Ala293 and Leu78 decreased the specificity of AgcA_EP4_ for alkylguaiacols. Substitution of Phe166 yielded a variant that bound 4-propylsyringol but did not transform it. In contrast, the corresponding variant in the *Rhodococcus aromaticivorans* RHA1 homolog, AgcA_RHA1_ Y166A, catalyzed the *O*-demethylated of both methoxy groups of 4-propylsyringol with a *k*_cat_*/K*_m_ of 8500 M^-1^ s^-1^ for the first *O*-demethylation, nearly 7-fold higher than WT AgcA_RHA1_. A strain of RHA1 harboring the variant did not grow on 4-propylsyringol but consumed it at approximately the same rate as 4-propylguaiacol and transformed some of it to pentanoyl-CoA, consistent with metabolism *via* the *meta*-cleavage pathway that catabolizes 4-alkylguaiacols. These studies improve our understanding of a critical lignin-degrading enzyme system and facilitate its efficient implementation into biocatalysts.

**Significance:** Lignin is a highly abundant source of aromatic carbon and a promising alternative to petroleum to generate materials. Fulfilling this promise depends on technological advances in areas such as catalytic fractionation and biocatalysis. Catalytic fractionation of hardwood biomass generates mixtures of aromatics enriched in 4-propylguaiacol and 4-propylsyringol. Here, we biochemically and structurally characterized a cytochrome P450 that initiates 4-propylguaiacol catabolism. Informed by the structure, we engineered the enzyme to have dual activity on both 4-propylguaiacol and 4-propylsyringol, and implemented this enzyme into a bacterial biocatalyst. Metabolomic analysis of this strain provided insights into the catabolism of both aromatics. Overall, these findings greatly facilitate the engineering of P450s and bacteria to biocatalytically upgrade lignin.

## Introduction

Lignin is an abundant, highly methoxylated aromatic biopolymer found in the woody biomass of vascular plants. The predominant building blocks of lignin are the phenylpropanoids *p*-coumaryl, coniferyl and sinapyl alcohol (1). These have zero, one and two methoxy groups, and form H-, G- and S-type units, respectively, in the lignin polymer. Lignin has been proposed as a feedstock to replace petroleum in the manufacturing of platform chemicals (2). One particularly promising strategy for valorizing lignin is a tandem chemo-biological process. In this workflow, biomass is first fractionated by chemo-catalysis to yield, among other products, a mixture of **l**ignin-**d**erived **a**romatic **c**ompounds (LDACs) (3, 4). This mixture is then biocatalytically converted by a microbial cell factory to the targeted platform chemical.

Importantly, the biocatalyst must be tailored to the LDAC mixture, as the chemical composition of the latter varies according to the type of biomass and the fractionation strategy. For example, reductive catalytic fractionation (RCF), one of several promising chemo-catalytic strategies, yields mixtures that contain high amounts of methoxylated 4-propylphenolics (5). The nature of the phenolics depends on the biomass: softwoods, whose lignin is rich in G-type units, yield predominantly 4-propylguaiacol (4PG) while hardwoods and grasses, whose lignin also contains S-type units, generate a mixture of 4PG and 4-propylsyringol (4PS) **(Figure 1A)**. Regardless of the proportion of 4PG and 4PS, *O*-demethylation remains a critical bottleneck in their biocatalytic conversion (6).

**Figure 1:**
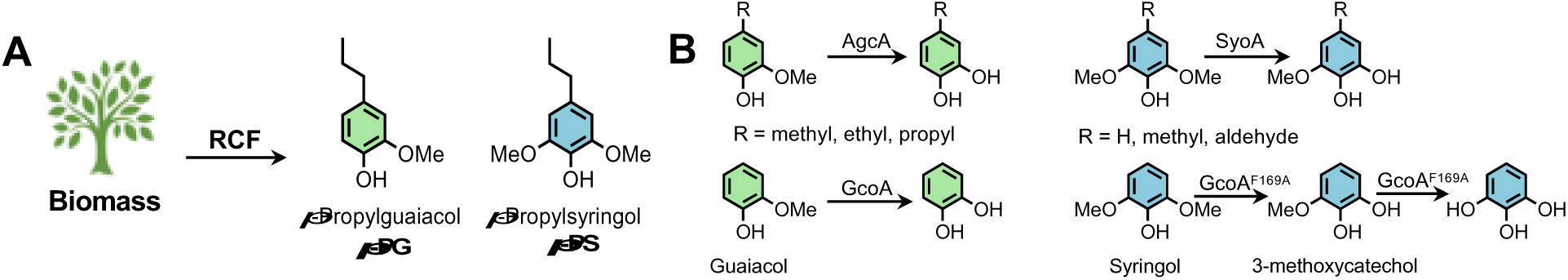
Summary of catalytic fractionation of biomass and key chemical structures and enzymatic reactions discussed in this study. (**A**) Reductive catalytic fractionation (RCF) of biomass generates the LDACs 4PG and 4PS in high amounts. (**B**) AgcA, GcoA and SyoA are cytochromes P450 that catalyze the *O*-demethylation of alkylguaiacols (7), guaiacol (8) and alkylsyringols (9), respectively. The F169A variant of GcoA *O*-demethylates syringol (10). For clarity, the co-substrates (O_2_, NADH) and the reductases (AgcB, GcoB) are not shown. G- and S-type LDACs are colored green and blue, respectively.

One class of enzymes that catalyze *O*-demethylation is cytochromes P450 (P450s), a superfamily of heme-thiolate enzymes that catalyze a wide variety of monooxygenation reactions (11). Two subfamilies of P450s have attracted interest for their abilities to transform LDACS: CYP199As and CYP255As are involved in the microbial catabolism of *para*-methoxybenzoates and guaiacols, respectively (12). Among CYP255As, AgcA and GcoA catalyze the *O*-demethylation of guaiacol and alkylguaiacols, respectively, to the corresponding (alkyl)catechol while SyoA catalyzes the *O*-demethylation of syringol to 3-methoxycatechol (3MC) **(Figure 1B)** (9). As is typical of P450-catalyzed reactions, *O*-demethylation requires reducing equivalents supplied by NAD(P)H in addition to O_2_. In the CYP255A enzymes characterized to date, a reductase, exemplified by GcoB, AgcB, and the predicted SyoB, transfer electrons from NADH to the P450 via a flavin and a [2Fe-2S] cluster.

Rhodococcal strains such as *Rhodococcus aromaticivorans* RHA1 (formerly *R*. *jostii* (13)) and *Rhodococcus rhodochrous* EP4, are able to grow on RCF mixtures (7). In these strains, AgcAB *O*-demethylates 4-alkylguaiacols to the corresponding 4-alkylcatechol. The latter is catabolized via the *meta*-cleavage Aph pathway, in which AphC catalyzes the extradiol ring cleavage (7, 14). The *meta*-cleavage product (MCP) is predicted to be catabolized to pyruvate and a fatty acyl-CoA whose chain length depends on the growth substrate. Thus, 4EG and 4PG are predicted to yield butanoyl- and pentanoyl-CoA, respectively. Interestingly, these strains also contain GcoAB, which initiates the catabolism of guaiacol (7). In contrast to 4-alkylcatechols, the resulting catechol undergoes intradiol ring cleavage and is catabolized *via* the β-ketoadipate pathway (14, 15). Importantly, these strains do not transform 4PS. Moreover, although SyoA catalyzes the *O*-demethylation of syringol, the subsequent catabolism of 3MC in its native host is unknown (9, 16). Finally, SyoA does not appear to tolerate significant substituents at the 4-position of its substrate (16). Overall, the apparent recalcitrance of 4PS to microbial transformation is a barrier for the valorization of this RCF product.

The ability of CYP255As to convert LDACs has motivated their further study and engineering. GcoAB_EP4_ and AgcAB_EP4_ from *R. rhodochrous* EP4 have complementary specificities with respect to alkyl chain length, with AgcA having highest specificity for 4PG and GcoA having highest specificity for guaiacol (7). The active site residues of GcoA_EP4_ are conserved in AgcA except for those thought to determine the respective specificities of these enzymes for 4-alkylguaiacols: Ile81 and Thr296 in GcoA correspond to a leucine and alanine in AgcA (**Figure S1**). The interest in the catabolism of S-lignin-derived units is such that GcoA_Amy_ of *Amycolaptosis* ATCC 39116 was engineered to transform syringol **(Figure 1B)** (10). More specifically, substitution of active site Phe169 with alanine generated a variant with high affinity for syringol, efficiently catalyzing its *O*-demethylation to 3MC **(Figure 1B)**. In contrast, GcoA F169A had a lower affinity for 3MC, and its *O*-demethylation was poorly coupled to NADH-and O_2_-utilization.

In this study, we used structural and biochemical approaches to examine critical functional aspects of AgcA_EP4_ and AgcA_RHA1_. Using structure-guided protein engineering, we then investigated the role of key residues in AgcA’s specificity for alkylguaiacols. We designed variants of the two P450s at position 166 to increase the activity for the underutilized monoaromatic 4PS and tested their activity on this substrate and 4PG. Finally, we evaluated the biocatalytic potential of AgcA and its variants by producing them in RHA1 and testing the ability of the resultant strains to degrade 4PS and 4PG. The results are discussed with respect to the engineering of lignin-valorizing enzymes and catabolic pathways.

## Results

### Activity of AgcAB_EP4_

Previously, the activities of AgcA_EP4_ and AgcA_RHA1_ were evaluated using a 30-fold excess of AgcB_EP4_ and a buffer comprising 20 mM MOPS, 90 mM NaCl, pH 7.2 (*I* = 100 mM) at 25 °C, (7). Importantly, AgcB_EP4_ efficiently reduced both homologs of AgcA and, due to its higher solubility, was easier to handle than AgcB_RHA1_. For the current studies, we optimized buffer conditions to maximize coupling between NADH consumption and *O*-demethylation. We found highest coupling efficiency and activity for both homologs using 10 mM MOPS, 20 mM NaCl (*I* = 25 mM), pH 7.2 (**Figure S2**) and therefore used this buffer in all subsequent experiments. Under these conditions, the activity of AgcA_EP4_ was approximately 3-fold higher than that of AgcA_RHA1_ (**Figure S2A**, **B**). Using the optimized buffer, AgcA had ∼65-fold higher affinity for 4PG than for guaiacol (**Table 1**). This is similar to what we observed at an ionic strength of 100 mM (7), although the *K*_d_ values were lower under the optimized conditions. Moreover, the P450 had ∼500-fold higher apparent specificity (*k*_ca*t*_*/K*_M_) for 4PG than for guaiacol. Again, this is similar to what we previously observed at an ionic strength of 100 mM (7).

**Table 1:**
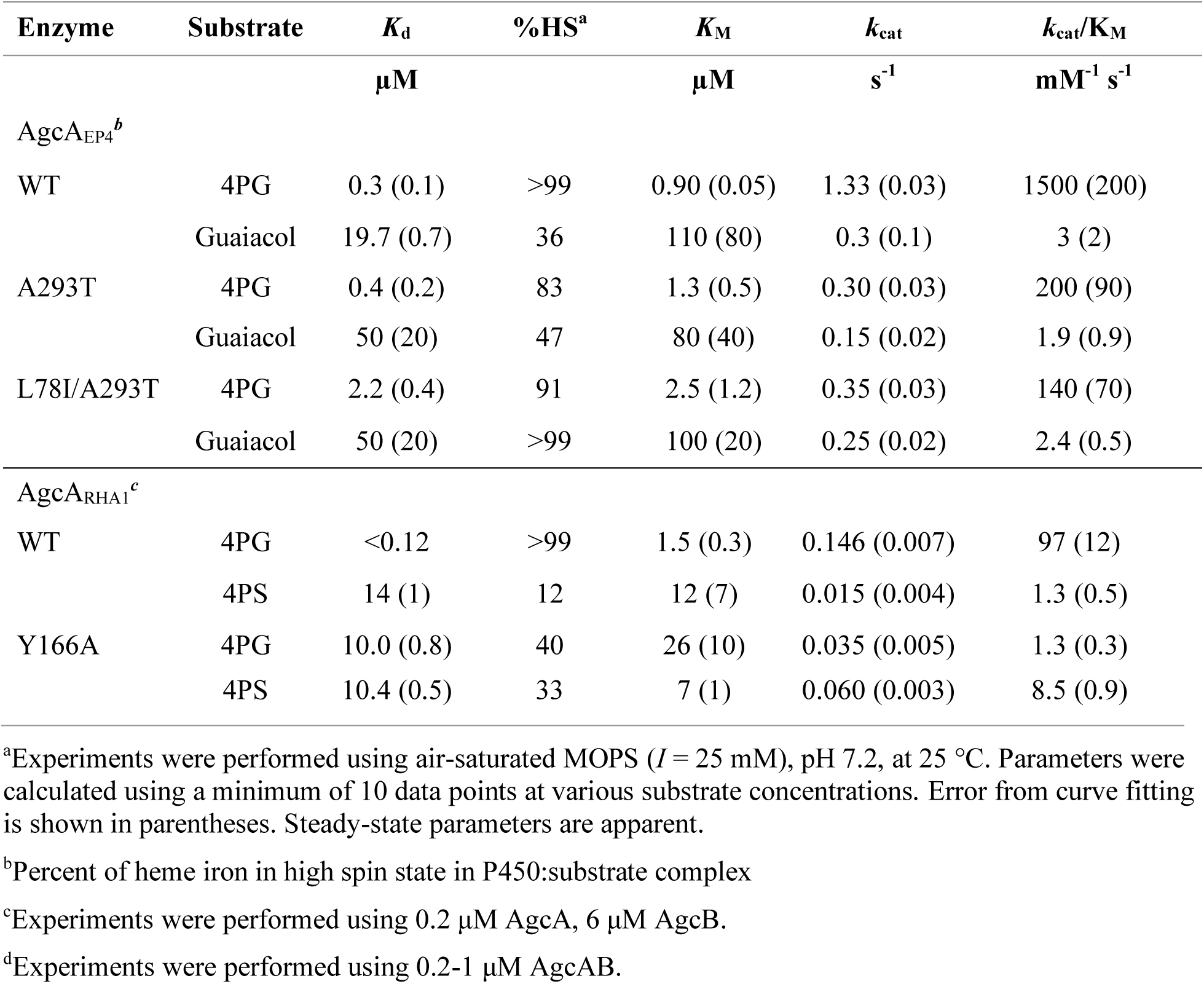
Binding and activity of AgcA and its variants on guaiacols and syringols^a^.

### The active site of AgcA_EP4_ reveals unique structural elements

To investigate the structural determinants of specificity, we solved the X-ray structure of AgcA_EP4_ in complex with 4EG to a resolution of 1.82 Å. Refinement data are summarized in **Table S1** The structure of AgcA is very similar to those of GcoA_Amy_ and SyoA (**Figure S3**), consistent with amino acid sequence identities of greater than 43%. For example, AgcA_EP4_ has an RMSD of 0.597 Å (C_α_) with respect to GcoA_Amy_. The 4EG substrate was modelled at full occupancy, with an average atomic B-factor of approximately 18 and 19 Å^2^, in chains A and B, respectively. These values are similar to those of the surrounding residues, indicative that the substrate is well-ordered within the active site. 4EG is orientated similarly to guaiacol in the active site of GcoA_Amy_ with comparable Fe-C distances between the heme iron and the methoxy carbon, (**Figure 2**). The alkyl side chain of the bound 4EG is accommodated by a pocket lined by Leu78 and Ala239. These correspond to Ile81 and Thr241 in GcoA_Amy_ and Leu85 and Ser300 in SyoA (**Figure 2A**). The reduced steric hindrance between the side chains of Leu78 and Ala239 with the alkyl chain of 4-alkylguaiacols may contribute to the increased specificity of AgcA for these compounds.

**Figure 2:**
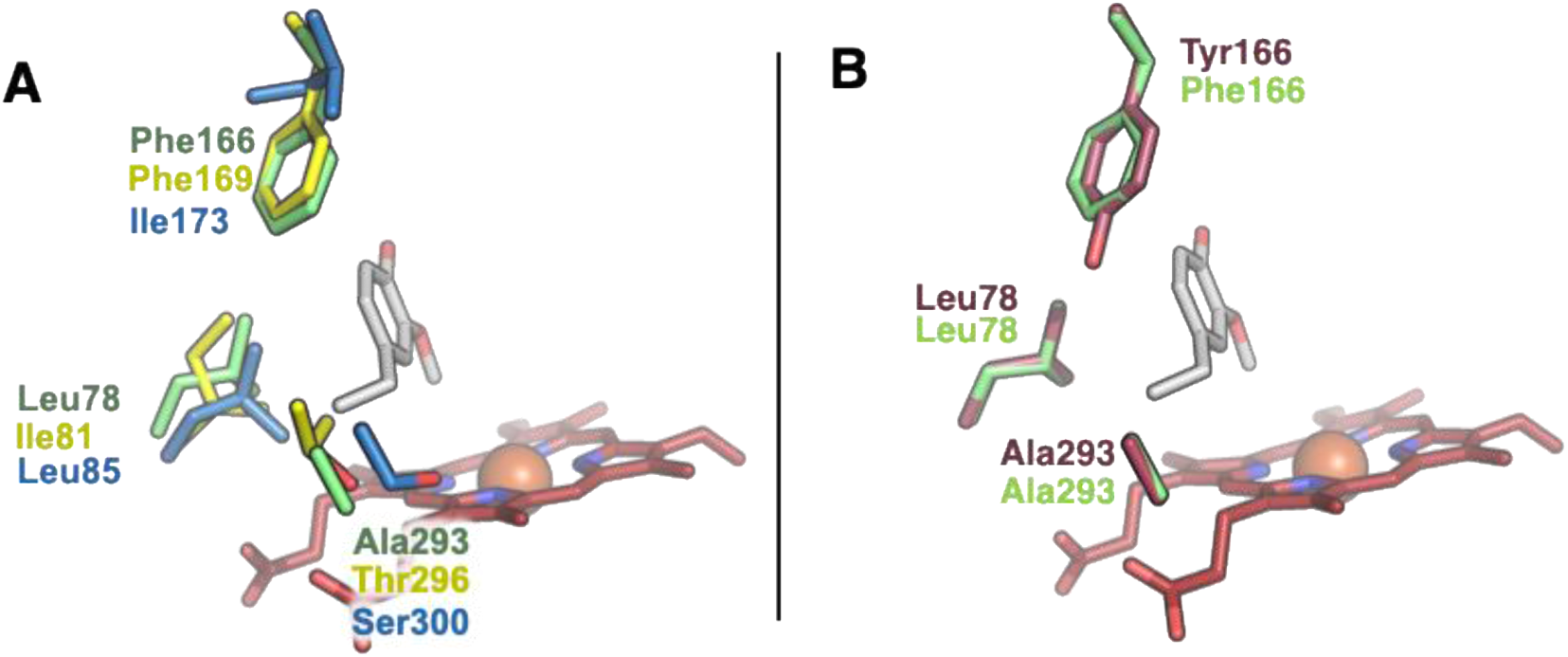
Structural features of AgcA and other CYP255As. (**A**) Comparison of key residues in the substrate binding pocket of AgcA_EP4_ (green), GcoA_Amy_ (yellow) and SyoA (blue). Structures shown are 9IA1 (AgcA), 5NCB (GcoA_Amy_) and 8U19 (SyoA). Fe_Heme_-C_methoxy_ distances are 3.8, 3.9 and 3.6 Å for Agc, GcoA_Amy_ and SyoA, respectively. (**B**) The corresponding residues in AgcA_RHA1_ (maroon) and AgcA_EP4_ (green) binding pocket.

To interrogate this hypothesis, we produced the A293T single and A293T/L78I double variants of AgcA_EP4_ and tested their affinity for the substrates 4PG and guaiacol (**Table 1**). Although substituting Ala293 for threonine did not significantly change the affinity of the enzyme for 4PG, the substrate did not induce the same extent of high spin conversion in the heme iron, suggesting that the substrate was less efficient at displacing the heme-coordinated water. Conversely, the double variant had an approximately seven-fold lower affinity for 4PG but a higher proportion of the substrate-bound enzyme was in the high-spin state. Interestingly, the variants had lower affinity for guaiacol, although guaiacol induced a higher proportion of high-spin iron in both variants than in the WT enzyme. To further investigate the relationship between these active site residues and enzyme function, we tested the activity of the variants on the same substrates. Consistent with the *K*_d_ values, the A293T and A293T/L78I variants had lower apparent specificity for 4PG but neither had significantly higher apparent specificity for guaiacol (**Table 1**).

### Activity of AgcA_EP4_ variants on 4PS

To date, there are no reported *O*-demethylases for 4PS, an LDAC that is generated in large amounts in the RCF of hardwoods and grasses. AgcA_EP4_ bound 4PS with reasonable affinity, but transformed it with a specific activity ∼1% that of 4PG (**Table S2**). We hypothesized that AgcA could be engineered to catalyze the *O*-demethylation of 4PS in a manner similar to how GcoA_Amy_ was engineered to transform syringol where Phe169 was substituted with alanine (10). Substitution of the corresponding residue in AgcA_EP4_, Phe166 (**Figure 2A**), for any one of four smaller residues of different polarities, yielded variants that bound 4PS with greater affinity than the WT P450 (**Table S2**) In addition, the four variants had lower affinity for 4PG than the WT P450. Of the five, the F166N variant bound 4PS with the greatest affinity (2 ± 1 µM) and, in this variant, 4PS induced the greatest spin state transition of the heme iron (45%). Unexpectedly, despite the relatively high affinity of the F166N variant for 4PS and 4PG, the variant’s specific activity for the two compounds was less than 1% that of WT AgcA_EP4_ for 4PG (**Table S2**). The specific activity of the F166A variant for 4PG and 4PS was similarly very low.

### Electron transfer to AgcA_EP4_ variants is disrupted

We investigated the basis of the low turnover of the Phe166 variants in AgcA_EP4_ F166N, the variant which had the highest affinity for both substrates. We used the strong CO-binding properties of the ferrous heme iron as a readout of reduction given the ease of measuring the shift of the Soret peak to 450 nm. In the presence of sodium dithionite, the F166N variant in complex with 4PS formed the CO-adduct **(Figure S4)**, establishing that the heme iron can be reduced by strong chemical reductants and is accessible to small gaseous ligands. Moreover, WT AgcA•4PG formed the CO-adduct when incubated with NADH and AgcB in CO-saturated buffer. By contrast, the F166N did not, indicating that AgcB is unable to reduce the heme iron of the variant effectively.

### Activity of AgcA_RHA1_ variants on 4PS

In light of the similar substrate specificities (7) and active site architectures (**Figure 2B**) of AgcA_EP4_ and AgcA_RHA1,_ we investigated whether the RHA1 homolog might be a better platform for designing a 4PS *O*-demethylase. Indeed, in preliminary experiments, AgcA_RHA1_ depleted 4PS at a rate of 0.8 min^-1^ µM heme^-1^, approximately an order of magnitude faster than AgcA_EP4_. Phe166 of AgcA_EP4_ corresponds to Tyr166 in AgcA_RHA1_ (**Figure 2B**). Incubation of the Y166A variant of AgcA_RHA1_ with 4PS yielded three new peaks, with *t*_R_ of 15.2, 16.2 and 17.1 min on HPLC (**Figure 3B**). LC-MS analysis of reaction mixtures detected putative features with the *m/z* of 3M5PC and 5PPG, the single and double *O*-demethylation products of 4PS (**Figure 3A**). Although authentic standards were not available for 3-methoxy-5-propylcatechol (3M5PC) and 5-propylpyrogallol (5PPG), their retention time shifts, *m*/*z* values, and additional data presented below suggest that the peaks at 17.1 and 16.2 min represent 3M5PC and 5PPG, respectively (**Figure 3**). The identity of the peak at 15.2 min is discussed below. By contrast, *O*-demethylation of acetosyringone, another S-type LDAC, was detected in only trace amounts by LC-MS in reactions containing either WT AgcA_RHA1_ or its Y166A variant (**Figure S5**). Interestingly, the Y166A variant also had significantly lower specific activity for acetovanillone than the WT enzyme (**Figure S5**).

**Figure 3:**
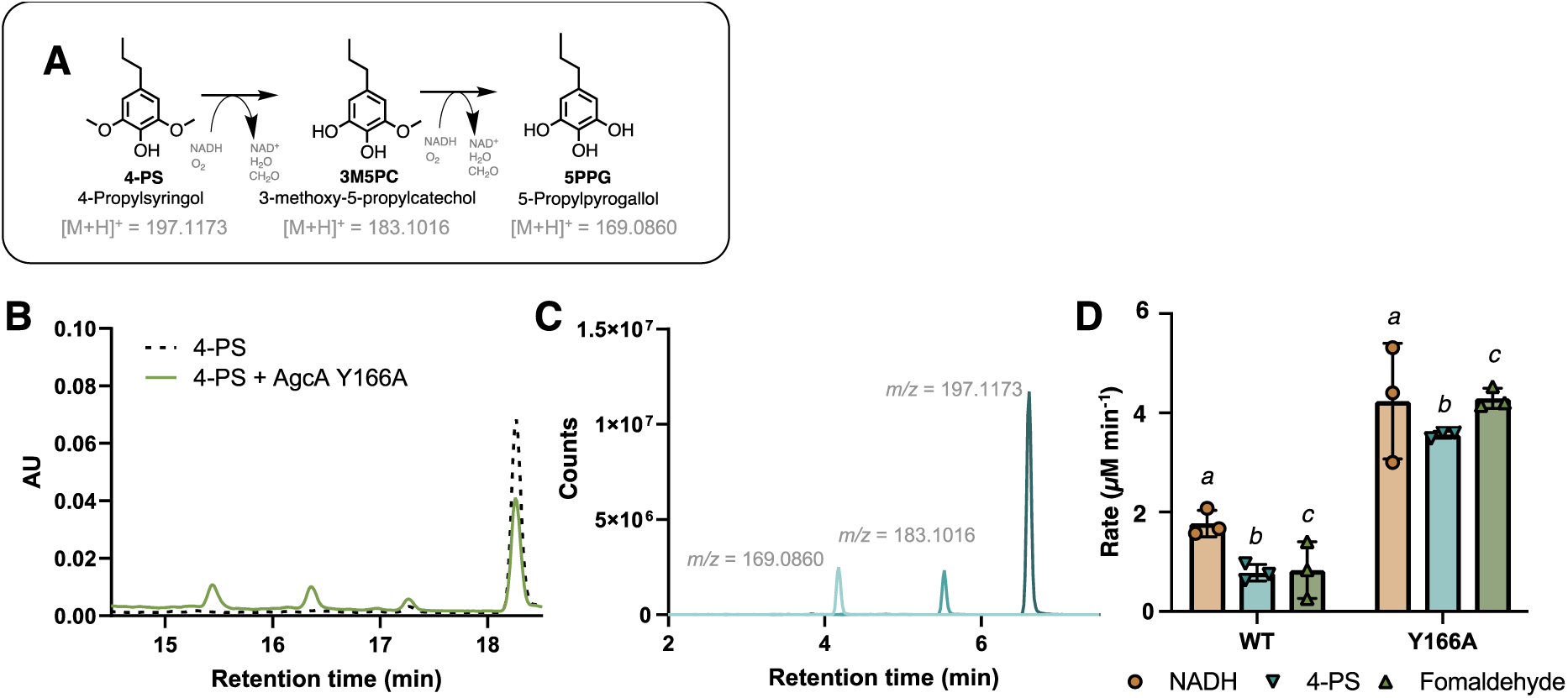
Transformation of 4PS by AgcA_RHA1_ Y166A: (**A**) Predicted O-demethylation scheme of 4PS, with putative products and predicted m/z of the protonated monoisotopic species. Reactions containing 1 µM AgcA_RHA1_ Y166A, 1 µM AgcB, 100 µM 4PS and 350 µM NADH in 10 mM MOPS, pH 7.2, I = 25 mM were incubated for 10 min at 25 °C, then quenched. (**B**) Chromatogram of the quenched reaction (green) and no enzyme control (dashed black). Absorbance at 270 nm. (**C**) LC-MS-ESI (+) analysis of the quenched reaction. Extracted ion chromatograms (EICs) of m/z corresponding to the substrate 4PS (dark blue) and predicted O-demethylation products, 3M5PC (aqua blue) and 5PPG (light blue). (**D**) Reaction rates of AgcA-catalyzed O-demethylation of 4PS. Rate of NADH and 4PS consumed, and formaldehyde calculated from 20 min (WT) or 10 min (Y166A) incubations. In addition to components listed above, reactions contained 1000 U/mL catalase. NADH, PS, and formaldehyde were quantified using spectroscopy, HPLC-UV, and a tryptophan-based colorimetric assay, respectively. For comparisons (a to a, b to b, and c to c), p < 0.0001.

To further confirm the identity of the 4PS transformation products, we took advantage of the commercial availability of 3-methylpyrogallol (3MPG), the double *O*-demethylation product of 4-methylsyringol (4MS) (**Figure S6A**). Although AgcA_RHA1_ has an approximately 20-fold lower specificity for 4-methylguaiacol *versus* 4PG (7), we tested the activity of the Y166A variant on methylsyringol. Reaction mixtures of the variant, AgcB, NADH and 4MS yielded a small amount of 3MPG (LC-MS *t*_R_ = 2.3 min), despite the instability of this compound in aqueous solution (17) (**Figure S6B**) as well as a product with an *m/z* corresponding to that of 3-methoxy-5-methylcatechol (3M5MC) (**Figure S6C**). Interestingly, reactions containing WT AgcA had a feature with an *m/z* corresponding to that of 3M5MC in similar quantities as the Y166A reaction (**Figure S6B**) but no detectable 5MPG (**Figure S6C**).

In investigating the identity of the third product yielded by the transformation of 4PS (LC-UV/Vis, *t*_R_ ∼ 15.1 min), we noticed that its abundance varied as a function of reactive oxygen species. Thus, higher concentrations of reductase (**Figure S7A**) or the addition of hydrogen peroxide (**Figure S7B**) increased its abundance, while lower concentrations of reductase (**Figure S7A**) and the addition of catalase (**Figure S7C**) lowered its abundance. LC-MS analysis revealed the presence of a prominent peak with the *m/z* = 181.09 (LC-MS *t*_R_ = 2.5 min), 1.4 min lower than putative 5PPG, 2.8 min lower than putative 5M3PC and 4.0 min lower than 4PS (**Figure S6D**). The protonated ion of *m/z* = 181.09 has the calculated mass of C_10_H_12_O_3_, corresponding to oxidized 3M5PC. Alkylated catechols can form *ortho*-quinones (*o*Q) and quinone methides (QM) in a process that may be catalyzed by P450s (18). Although they have short half-lives, QMs may rearrange to form isomers such as alkenylsyringol derivatives (**Figure S7E**) (19). For their part, quinones are trapped as adducts by strong nucleophiles such as glutathione (GSH) (20). To better understand the reactivity of 4PS, we incubated it with 10 mM H_2_O_2_ and 20 mM GSH. This reaction yielded three GS-adduct peaks, the distribution of which depended on the presence of AgcA_RHA1_ Y166A. Incubations of AgcA Y166A, 4PS, H_2_O_2_ and GSH resulted in the formation of a GS-quinone adduct (**Figure S8**). MS/MS fragmentation of the GS-conjugates suggests a mix of benzyl and aromatic adducts, which form from the QM and *o*Q, respectively (21). Based on these experiments, we concluded that the compound at 15.2 min is a stable form of oxidized 5M3PC formed through a quinone intermediate, though the structure of this compound is not known. We also added catalase to all further reactions to prevent the side reaction of 4PS and its derivatives.

### Binding and activity of AgcA_RHA1_ Y166A variant

We next investigated the affinity and substrate specificity of AgcA_RHA1_ Y166A for 4PG and 4PS. The variant had similar affinities for 4PG and 4PS, and these compounds induced the same extent of spin transition in the heme iron. This affinity was two orders of magnitude lower than WT AgcA_RHA1_ for 4PG but similar in magnitude to the WT enzyme’s affinity for 4PS (**Table 1**). However, the latter induced considerably lesser spin-state conversion in WT AgcA_RHA1_. We next evaluated the coupling of the AgcA_RHA1_-catalyzed *O*-demethylation of 4PS by monitoring three parameters: 4PS and NADH consumption and formaldehyde production. In reactions with the variant, significantly higher amounts of each analyte were consumed (for NADH and 4PS) or generated (for formaldehyde) as compared to reactions with WT (**Figure 3C, Table S3**), consistent with the higher activity of the variant. For example, the yield of formaldehyde in 10 min-reactions containing Y166A was over five-fold higher than that of 20-min reactions containing WT AgcA_RHA1_ Moreover, in reactions with AgcA_RHA1_ Y166A, the amount of formaldehyde produced corresponded remarkably well to the amount of NADH oxidized, indicating that the *O*-demethylation is essentially 100% coupled in the variant. By contrast, in reactions with WT AgcA_RHA1_, the amount of formaldehyde produced corresponded to approximately 45% the amount of NADH oxidized, indicating that the reaction is significantly uncoupled. In these same reactions with WT AgcA_RHA1_, the amount of 4PS depleted corresponded well with the amount of formaldehyde detected, suggesting that the enzyme only catalyzed the first *O*-demethylation of 4PS over the course of 20 min. Consistent with this interpretation, only the peak provisionally assigned to 5M3PC (*t*_R_ = 17.2 min) was detected in HPLC analysis. By comparison, in reactions with the variant, less 4PS was depleted than NADH consumed or formaldehyde produced, consistent with the detection of 5PPG. Together, these results indicate that Y166A catalyzes both *O*-demethylation steps within 10 min and both are well-coupled to NADH depletion.

The coupling analysis combined with the binding data indicate that the amino acid substitution enables the optimum orientation of 4PS in the binding pocket. To further investigate the activity of the enzyme, we measured the apparent steady-state kinetics of *O*-demethylation by AgcA WT and Y166A for 4PS and 4PG. The activity of both enzymes on both substrates obeyed Michaelis-Menten kinetics (**Figure S9**). The apparent specificity of AgcA_RHA1_ Y166A for 4PS was 6.5-fold higher than for 4PG and 6.8-fold higher than WT AgcA_RHA1_’s apparent specificity for 4PS, representing a significant gain-of-function (**Table 1**). Nevertheless, the apparent specificity of the variant for 4PS was approximately an order of magnitude lower than that of the WT enzyme for 4PG.

### Activity of AphC_RHA1_ on demethylation products of 4PS

In RHA1 and EP4, AphC efficiently cleaves the catechols resulting from the *O*-demethylation of alkylguaiacols by AgcAB (22). As these catechols differ from the *O*-demethylation products of 4PS by a single methoxy group, we evaluated the ability of AphC to cleave 3M5PC and 5PPG (**Figure S10A**). HPLC analysis of reactions of AgcA_RHA1_ Y166A and 4PS indicates that the inclusion of 15 µM AphC resulted in the depletion of 3M5PC and the oxidized 3M5PC derivative (*i.e*., the peaks with *t*_R_ = 17.1, and 15.2 min) and the appearance of compound with a broad peak with a *t*_R_ of 14.6 min (**Figure S10B**). Features with *m*/*z* values corresponding to the MCP of 3M5PC ([M-H]^-^ *m*/*z* = 213.08) and 5PPG ([M-H]^-^ *m*/*z* = 199.06) were detected *via* LC*-*MS in the above reaction (**Figure S10C**). No such products were detected in the AphC-free control (**Figure S10D**).

### Transformation of 4PS by engineered RHA1

Given the abilities of AgcA_RHA1_, its Y166A variant, and AphC_RHA1_ to transform 4PS and its catechols, we tested the ability of RHA1 to grow on and transform 4PS. Using the pRIME, we integrated genes encoding AgcA_RHA1_ WT and Y166A into RHA1 under the control of the constitutive T_1_ promoter (23). In whole cell turnover assays, the strain overexpressing the variant, RHAMW32, depleted 4PS and 4PG at similar rates **(Figure 4, Table S4)**. Strains overexpressing either the WT enzyme or the empty vector (RHAMW31 and RHAMW30, respectively) turned over 4PG at a similar rate as RHAMW32 **(Table S4),** although at t = 60 min, supernatants of cultures of RHAMW31 contained significantly less 4PG (**Figure 4A**). On the other hand, neither strains RHAMW31 nor RHAMW30 depleted 4PS at rates significantly faster than baseline **(Figure 4B**, **Table S4)**.

**Figure 4:**
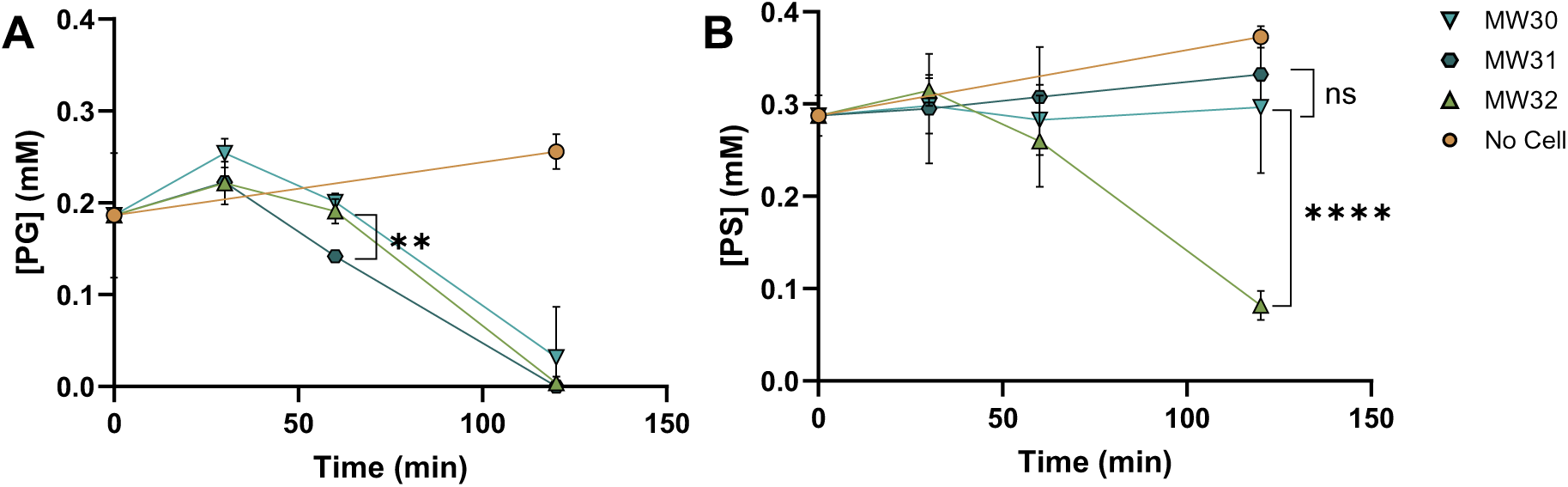
Whole cell turnover of 4PG and 4PS by engineered RHA1. Depletion of (**A**) 0.2 mM 4PG and (**B**) 0.3 mM 4PS by: RHAMW30, blue downward triangles; RHAMW31, teal hexagons; RHAMW32, green upward triangles; and cell-free incubations, orange circles. Cells were incubated in 500 mL M9G, 0.2% glucose, and the appropriate aromatic shaking at 30 °C. Values are the average of biological triplicates. Error bars represent standard deviation. Statistical comparison in panel **A**, *p* = 0.0026, panel **B**, *p* < 0.0001.

### Induction of meta-cleavage activity by 4PS

The ability of strain RHAMW32 to transform 4PS indicates that this compound induces *agcAB* in RHA1. More specifically, only the variant *agcA* is constitutively expressed in RHAMW32: the requisite reductase, AgcB, must be induced in sufficient amounts to drive 4PS *O*-demethylation. To investigate the induction of the downstream *aph* genes, we measured the *meta*-cleavage activity of AphC using 4-methylcatechol (4MC), a good substrate for the enzyme, in RHAMW32 cells incubated with 4PS and 4PG. After ∼24 h, clarified lysates of cells incubated with 1 mM 4PS had similar *meta*-cleavage activity as lysates of cells incubated with 4PG (**Table S4**). These activities were ∼50-fold higher than in lysates of cells incubated with glucose alone.

### Growth of strain RHAMW32 on 4PS

Strain RHAMW32 did not detectably grow on 4PS as detected *via* either an increase in OD_600_ **(Figure S11A)** or CFU/mL **(Figure S11B)** despite the fact that this strain efficiently depleted 4PS and that *meta*-cleavage activity was induced (**Table S5**). Interestingly, 0.5 mM 4PS inhibited the growth of RHAMW32 but not RHAMW31 on acetate, suggesting that a 4PS metabolite is cytotoxic. Finally, the culture supernatants of RHAMW32 cells turned brown when incubated with 4PS **(Figure S11C)**, suggesting the accumulation of oxidized catechols.

### Transformation of 4PG and 4PS by strain RHAMW32

We hypothesized that 4PS is metabolized by the Aph pathway, which is responsible for 4-alkylguaiacol catabolism via *meta*-cleavage (**Figure 5A**). To test this hypothesis, we first sought to further validate the pathway as only the first two steps have been experimentally verified (7, 22). More specifically, the six subsequent steps are based on bioinformatic predictions and do not account for two genes of unassigned function in the *aph* gene cluster: *nfox* and a homolog of *aphE*. For this experiment, we incubated concentrated cells of RHAMW32 with 0.5 mM 4PG or 4EG for 2 h and analyzed the culture supernatant and cell extracts for the predicted metabolites. In the 4PG-incubated samples, Features 1a and 2a, corresponding to 4PG and 4-propylcatechol (4PC), respectively, had low peak areas in the culture supernatants, consistent with these compounds being largely catabolized (**Figure 5B**). However, features with the expected *m*/*z* values of six of the other eight predicted metabolites were detected in significant amounts (**Table S6**). Feature 7a corresponds to pyruvate, as validated with comparison to an authentic standard. Similarly, a feature corresponding to pentanoyl-CoA (Pe-CoA) was detected in cell extracts (**Figure S12**). The other features were provisionally assigned to metabolites based on *m*/*z* values. No feature was detected that corresponded to pentanal, the predicted product of AphH and the substrate of AphI. Based on the homology of these enzymes with HsaFG, pentanal is predicted to be channeled between the active sites of AphHI without being released to the bulk solvent (24).

**Figure 5:**
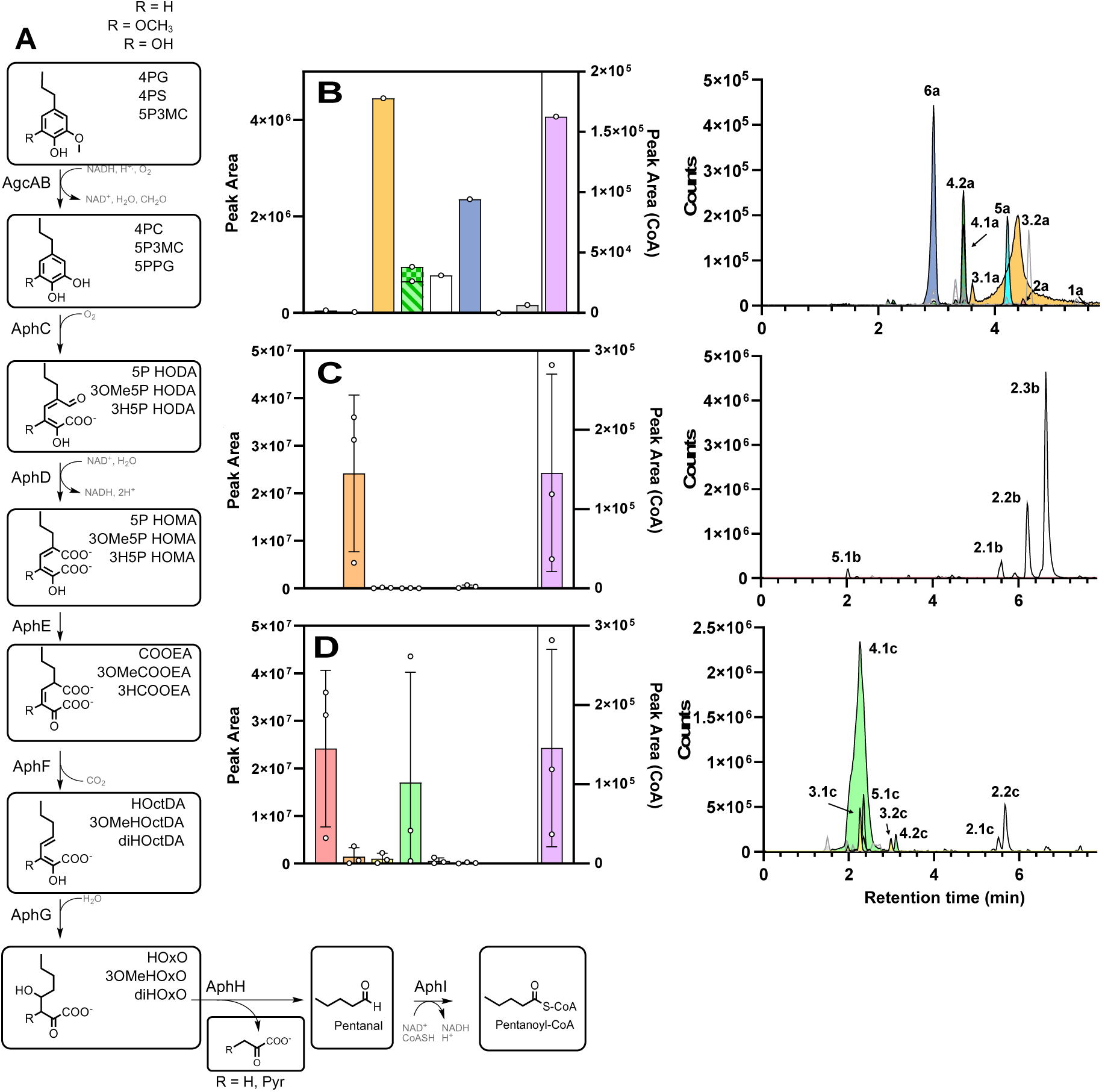
Transformation of 4PG and 4PS by RHA1. (**A**) 4PG and 4PS metabolites predicted to be generated by the Agc-Aph pathway. Area under the curve and EICs of features detected in culture supernatants with *m/z* values of predicted 4PG (**B**) and 4PS (**C**, **D**) metabolites (negative mode). Pentanoyl-CoA (Pe-CoA) was detected in cell extracts (positive mode). Dashed, checkered, and solid bars indicate the area under the EIC corresponding to the predicted *m*/*z* values of [M-H_2_O-H]^-^, [M-CO_2_-H_2_O-H]^-^ and [M-H]^-^, respectively. The identities of pyruvate and Pe-CoA were verified using authentic standards. The other numbered features were provisionally assigned to metabolites based on *m/z* values (**Table S6**). Error bars represent the average of three biological variants, where appropriate.

Among the features provisionally assigned to 4PG metabolites, the most prominent have *m*/*z* values corresponding to 2-hydroxy-6-oxo-5-propylhexa-2,4-dienoate (5P HODA) and 4-hydroxy-2-oxooctanoate (HOxO) (**Figure 5B**). Indeed, two features had *m*/*z* values corresponding to 5P HODA, 3.1a and 3.2a, which is consistent with what we have observed for other MCPs and likely represent isomers (25). Features 4.1a and 4.2a correspond to dehydrated and dehydrated/decarboxylated forms of the isomeric metabolites 5P HOMA and COOEA, respectively. We assigned these features to the dienolate 5P HOMA as it is predicted to be more stable. Moreover, enols are readily dehydrated in acidic solution, such as the LC/MS solvent, and, in electrospray ionization, decarboxylation can occur in-source (26). Hydroxy-octadienoate (HOctDA), the predicted product of AphG, is an alkylated analog of 2-hydroxypent-2,4-dienoate, the product of XylJ and NahK, which is reactive and prone to rapid tautomerization in aqueous solution (26). We have provisionally assigned Feature 5a to the more stable, keto form of this compound.

Samples from 4EG-incubated cells did not contain any of the features described above except for Features 4.1a and 6a, which were found in trace amounts. Pyruvate, which is also predicted to be a metabolite of 4EG, was detected similarly to 4PG-incubated cells (**Figure S13**). However, these samples contained features corresponding to the same metabolites less a methylene group (-CH_2_-), consistent with the shorter side chain of 4EG (**Figure S13B, C**). Interestingly, the features corresponding to the initial pathway metabolites had stronger signals than in 4PG-incubated samples. Finally, butanoyl-CoA (Bu-CoA) was detected in 4EG-incubated cells and Pe-CoA was not (**Figure S12A**), consistent with the predicted catabolism of 4EG by the Aph pathway. Overall, the metabolic profiles of the 4PG- and 4EG-grown cells support the annotation of the Aph pathway enzymes.

### Transformation of 4PS by strain RHAMW32

Having better validated the Aph pathway, we next examined the extent of 4PS catabolism by RHAMW32. To maximize 4PS transformation, we incubated cells with 4EG for 4 h to induce the Aph pathway, washed them, transferred them to media containing 10 mM glucose plus either 1 mM 4PG or 4PS, and incubated them for 6 h. We analyzed culture supernatants and cell extracts in essentially the manner described above. In 4PG-incubated cells, neither 4PG nor 4PC were detected, consistent with the longer incubation time, and we observed a significant accumulation of HOxO based on the intensities of Feature 6.1a (**Figure S14**). By contrast, the supernatants of 4PS-incubated cultures contained very prominent features corresponding to 5-propyl-3-methoxycatechol (2b) (**Figure 5C**) and 3H5P HOMA/3HCOOEA (4.1c) (**Figure 5D**). We also observed multiple features with *m*/*z* values corresponding to 3OMe5P HODA (3.1b, 3.2b) **(Figure S15A)** and 3H5P HODA (3.1c, 3.2c) (**Figure S15B**) consistent with MCP isomers (25). Features corresponding to six of the other ten predicted metabolites were also detected in the culture supernatant (**Figure 5C, D**). Interestingly, many of the predicted 4PS metabolites ran as multiple peaks in the EIC of the [M-H]^-^ ion. For example, the EIC of the predicted *m/z* of deprotonated 3OMe HOctDA and diHOctDA had three features each (5.1b, 5.2b, 5.3b and 5.1c, 5.2c, and 5.3c, respectively). However, none of them ionized with the neutral loss of water, in contrast with several 4PG metabolites. Finally, the predicted end-product of the pathway, Pe-CoA, was detected in extracts of 4PS-incubated cells (**Figure 5C, D**). A small amount of Bu-CoA was also detected, presumably originating from the 4EG used to induce the *aph* genes, as was acetyl-CoA, a central metabolite (**Figure S12E**).

## Discussion

The structural and functional characterization of the AgcAs and their variants as well as the engineered strains informs the first steps in the valorization of 4PS, which is generated in high yields but for which no biological transformations have been described. The AgcA homologs from RHA1 and EP4 efficiently catalyze the *O*-demethylation of alkylguaiacols. Consistent with the plasticity of P450s, we used structure-guided engineering to greatly increase the specificity of AgcA_RHA1_ for S-type units. Surprisingly, corresponding variants of AgcA_EP4_ were largely inactive on both G- and S-type alkylated LDACs. A strain of RHA1 expressing the engineered AgcA_RHA1_ consumed 4PS, unlike strains expressing the WT enzyme. Metabolomic analysis of the engineered strain further validated the Agc/Aph pathway and established that although the strain was unable to grow on 4PS, a proportion of it was catabolized to central metabolites.

The X-ray crystal structure of AgcA_EP4_ is highly similar to that of GcoA and revealed features responsible for the different specificities of CYP255As. Comparison of the structures of AgcA and GcoA identified Ala293 and Leu78 in AgcA as likely determinants of specificity (*k*_cat_/*K*_m_) for 4PG versus guaiacol (**Table 1**). Consistent with this analysis, the A293T variant and the A293T/L78I double variant had significantly reduced specificity for 4PG. Nevertheless, these substitutions did not increase AgcA’s specificity for guaiacol. The role of these residues as specificity determinants is supported by the T296A variant of GcoA, which had almost an order of magnitude less activity on guaiacol than the WT enzyme (27). Although the specificity of this variant for alkylguaiacols is not known, the low specificity of the AgcA variants indicates that residues in addition to those lining the substrate-binding pocket significantly influence the specificity of AgcA and GcoA for their respective guaiacols.

Working with the two AgcA homologs revealed a fascinating difference in their amenability to engineering. Thus, the structural data identified a steric clash between the aromatic residue at position 166 and the additional methoxy group of 4PS *vis-à-vis* 4PG (**Figure 2**). Consistent with this analysis, replacement of this residue with the smaller alanine in both AgcA_RHA1_ and AgcA_EP4_ yielded variants with higher affinity for 4PS. In accordance with this affinity, AgcA_RHA1_ Y166A efficiently *O*-demethylated both 4PG and 4PS, with the latter reaction being well-coupled to NADH and O_2_ consumption (**Figure 3D**, **Table S3**). In contrast, AgcA_EP4_ F166A was largely inactive on both 4PS and 4PG. Experiments using CO to trap the reduced iron revealed that the reductase reduced the WT enzyme but not the F166N variant in the presence of 4PG (**Figure S4**). In this state, it is unclear whether 4PS is productively positioned for catalysis. Interestingly, Phe166 is located on the F-helix which, along with the G-helix, shifts upon substrate binding to modulate the entrance of the substrate access channel of P450s (28). Further structural studies are required to determine whether substitution of Phe166 disrupts how the P450 binds the aromatic substrate. Other than Phe166/Tyr166, the residues forming the substrate-binding pocket are conserved between the RHA1 and EP4 homologs (**Figure S16**). It therefore appears that the starkly different activities of the variants is due to non-conserved features in other regions of the enzyme. Regardless, this difference between the homologs underscores the utility of exploring diverse enzymes when engineering biocatalysts.

The metabolic profiles of the engineered strain RHA1MW32 incubated with 4PG and 4EG are consistent with the proposed Aph pathway. Detection of the proposed pathway end-products, pyruvate and the acyl-CoAs (Pe- and Bu-CoA for 4PG and 4EG, respectively) (**Figures 5** and **S12-14**) validate the last step of the pathway and provide the first direct evidence for steps downstream of the AphC-catalyzed *meta*-cleavage of the alkyl-catechol (7). Moreover, features detected in culture supernatants were provisionally assigned to four of the other five pathway metabolites based on the homology of the Aph enzymes to the Dmp, Nah and Xyl enzymes, which transform catechols in the respective dimethylphenol, naphthalene and xylene catabolic pathways (29–31). More specifically, based on their *m*/*z* values, we provisionally assigned features to the MCP, hydroxymuconate, hydroxydienoate and hydroxyoxonoate. These assignments are supported by the respective features having similar retention times and peak distributions in the two samples yet differing in *m*/*z* values by an amount corresponding to the different alkyl chains of the two substrates. Nevertheless, these data provide no insight into the two genes in the *aph* gene cluster of unassigned function. The first of these, *nfor*, encodes a YjgF/Rid-family enzyme, and occurs in other gene clusters encoding *meta*-cleavage pathways (32, 33). The other, encodes a predicted two-domain isomerase, one domain of which is homologous to AphE. This enzyme may catalyze an additional tautomerization step in the Aph pathway. Gene deletion studies and/or characterization of the enzymes is necessary to further validate the pathway.

Although an RHA1 strain overproducing AgcA_RHA1_ Y166A did not grow on 4PS, it depleted this compound at approximately the same rate as 4PG. Analysis of supernatants of these cells showed a large accumulation of 5P3MC (**Figure 5**), suggesting that AgcA_RHA1_ Y166A has lower specificity for this methoxycatechol than for 4PS. Moreover, the apparent low abundance of methoxylated downstream metabolites suggests that 5P3MC is a poor substrate for AphC despite the enzyme’s ability to cleave 5P3MC *in vitro* and induction of considerable *meta*-cleavage activity by 4PS or a catabolite thereof (**Table S5** and **Figure S10**). Modelling 5P3MC into the binding pocket of AphC revealed a significant steric clash between the 3-methoxy O of 5P3MC and the O_β_ of Thr273, which lines the substrate-binding pocket (**Figure S17**). More specifically, the predicted O-O interatomic distance is 1.9 Å. Combined, these data suggest the specificity of AphC is a bottleneck for the catabolism of the methoxylated species.

The relative abundance of hydroxylated metabolites in 4PS-incubated cells indicates that 4PS is primarily converted by this branch and identifies potential bottlenecks. More specifically, the accumulation of a compound with the predicted *m/z* value of the tautomers 3H5P HOMA and 3H COOEA (**Figure 5**) suggests that AphE is a bottleneck. AphE is a member of the 4-oxalocrotonate tautomerase (4-OT) family (36) and is predicted to convert the more stable enol, 3H5P HOMA to its keto form, 3H COOEA. The latter is proposed to be decarboxylated by AphF, which shares approximately 58% amino acid sequence identity with the Mg^2+^-dependent decarboxylases DmpH, NahK, and XylI. Importantly, these enzymes are specific for the keto form of 4-oxalocrotonate (34, 35). The proposed mechanism of the tautomerase involves a proton transfer from the α-hydroxyl of the enol to δ-carbon of the substrate *via* an N-terminal-proline. Substitution of Pro1 with alanine or glycine resulted in a 70-fold reduction in the *k*_cat_ value of 4-OT (37). Based on the proposed mechanism, the β-hydroxyl in 3H5P HOMA would be expected to severely impair access of the catalytic proline of AphE to the substrate, reducing the activity of the tautomerase (**Figure S18**). Overall, the abundance of Feature 4.1c is consistent with inhibition of AphE and the accumulation of 3H5P HOMA.

Despite the apparent bottlenecks in 4PS catabolism, the accumulation of Pe-CoA, the predicted end product of this pathway, in amounts comparable to cells incubated with 4PG (**Figure 5**), indicates that a significant amount of 4PS is catabolized by the Aph pathway. At the same time, RHAMW32 did not grow in the presence of 4-PS (**Figure S11B**), in contrast to RHAMW30 and RHAMW31, which are unable to transform 4PS. This strongly suggests that one or more of the accumulated metabolites inhibits growth. This toxicity could be due in part to 5P3MC. Catechols are cytotoxic (38) and the accumulation of 5P3MC is indicated by both the LC-MS data and the brown coloration of the spent media.

Our study presents a number of opportunities for valorizing 4PS, an underutilized LDAC. First, AgcA Y166A facilitates the formation of quinones from the *O*-demethylated product of 4PS in the presence of H_2_O_2_. Glutathione trapping studies strongly suggest the formation of a *para*-QM **(Figure S7)**. QMs are emerging as building blocks in the synthesis of a variety of molecules and materials (39), and their formation from an abundant LDAC could unlock new applications. At the same time, QMs can be deleterious to biological systems, rendering this reactivity an important consideration for biocatalysis (18). Second, the methoxycatechol that accumulates in the culture supernatants of 4PS-incubated cells of RHA1 engineered with AgcA Y166A belongs to a recently identified class of valuable compounds for new polymer chemistry (40). Finally, the development of a strain that can convert both 4PS and 4PG is a critical first step in developing a microbial cell factory for valorizing these LDACs through biological funneling. Although there are bottlenecks in 4PS catabolism by RHAMW32, their identification facilitates metabolic optimization through rational engineering and/or adapted laboratory evolution. Overall, these new insights inform the generation of novel biocatalysts to convert alkylated S-type lignin units.

## Materials and Methods

Further details are provided in the SI Appendix

### Chemicals and Reagents

Reagents were of analytical grade and were obtained from Sigma-Aldrich and Fisher Scientific. Pentanoyl- and butanoyl-CoAs were supplied by Cayman Chemicals. 4PS was provided by Dr. Gregg T. Beckham at the National Laboratory of the Rockies (Golden, CO), synthesized as described by Brandner *et al.* (41). Media were prepared using water purified on a Barnstead NANOpure UV apparatus to a resistivity of greater than 16 MΩ/cm.

### DNA manipulation and strain construction

Bacterial strains and plasmids used in this study are listed in **Tables S7** and **S8**, respectively. DNA was propagated, purified, and manipulated using standard procedures (42). The *agcA* gene fragments containing point mutations were synthesized from TWIST Biosciences (San Francisco, USA) with appropriate overlaps for cloning into the pET15b plasmid using Gibson assembly (**Table S8**). Engineered *Rhodococcus* strains (RHAMW30, RHAMW31 and RHAMW32) were constructed by transforming Gibson-type assemblies composed of digested pRIME vector, the appropriate amplified insert (**Table S9)** and NEBuilder^®^ HiFi DNA Assembly Master Mix (New England Biolabs) into RHA1. Insertion was validated by PCR screening and sequencing.

### Protein production, purification and quantification

AgcA and its variants were produced heterologously *in E. coli* as N-terminally polyHis-tagged (Ht-) proteins and isolated using immobilized metal affinity chromatography (IMAC), followed by TEV protease-mediated Ht-cleavage, heme reconstitution and ion-exchange chromatography. Ht-AgcB was produced heterologously in *E. coli* and isolated by IMAC, conducted anaerobically. AgcA heme concentration was calculated by a carbon monoxide binding assay. AgcB FAD concentration was determined by absorbance at 450 nm, ε = 11.3 mM^-1^ cm^-1^. For enzyme activity assays, the concentrations of AgcA and AgcB were defined by the concentrations of heme and FAD, respectively.

### Protein analysis

Binding constants were determined by titration of substrates into solutions of ∼1 µM AgcA. Spectra were recorded after addition of substrate and subtracted from the spectrum of the resting-state enzyme. The absorbance difference between the peak and trough of the subtracted spectra was plotted against ligand concentration and *K*_d_ values were determined by fitting the quadratic equation. Activity assays for coupling were conducted using 1 µM of AgcA and AgcB in 200 µL 10 mM MOPS, pH 7.2 (*I* = 25 mM), 25 °C. Ionic strength was adjusted using NaCl. The reaction was initiated by adding NADH to 350 µM. NADH concentration was measured by absorbance at 340 nm, ε = 6.22 mM^-1^ cm^-1^. Coupling was calculated from the ratio of rate of aromatic turnover, as determined by HPLC, and the rate of NADH oxidation, determined spectrophotometrically. Formaldehyde was determined *via* a tryptophan-based spectrophotometric assays as described previously (8). HPLC, spectrophotometric and colorimetric assays are detailed in the SI Appendix. Apparent steady-state kinetics were determined using a coupled assay with AphC or XylE, as described previously (7), for AgcA_EP4_. For AgcA_RHA1_, NADH depletion rate was measured spectroscopically (ε = 6.22 mM^-1^ cm^-1^) and initial rates were appropriately adjusted by coupling to yield the estimated aromatic turnover rate. Apparent steady-state kinetic parameters were calculated by fitting Michaelis-Menten equations to initial velocity of reactions at various concentrations of aromatic substrate (LEONORA).

### LC-MS analysis

Enzyme reactions were analyzed by C18-LC-MS and LC-MS^2^. Pathway metabolites, cell extracts and spent media were analyzed HILIC-LC-MS. Full system parameters are in SI Appendix.

### Whole cell and growth assays

For the growth experiment, single colonies of RHAMW30, RHAMW31 and RHAMW32 were cultured in 2 mL of Luria Broth (LB), supplemented with 0.5 4PG. At OD_600_ ∼0.5, cells were washed 2× in M9 minimal media and resuspended in 500 µL M9 minimal media supplemented with minerals (M9G) with the appropriate substrate (Ref for M9G). Cells were incubated in a 48-well plate for 42 hours at 30 °C shaking at 200 rpm. OD_600_ was measured with a TECAN Spark. For the whole-cell turnover experiment, single colonies of RHAMW30, RHAMW31 and RHAMW32 were cultured in 2 mL of Luria Broth (LB) and used to inoculate 100 mL LB. Cells were collected when OD_600_ reached ∼3, washed twice in M9 and suspended in 500 µL M9G with 0.2% glucose and 0.2 mM of 4PG or 0.3 mM 4PS at an OD_600_ of 10 in a 48-well plate. The remaining substrate was quantified by HPLC. To detect products of the Agc/Aph pathway in RHAMW32, a single colony of RHAMW32 was cultured in 2 mL LB. This culture was used to inoculate 50 mL of M9G with 0.5 mM 4-EG. Cells were collected at an OD_600_ of 0.2, washed 2× in M9 and concentrated to an OD_600_ of 1. Five mL aliquots of the concentrated culture were incubated with 0.2% glucose and 0.5 mM of 4PS or 4PG for 3 h at 30 °C, at which point cells were harvested by centrifugation (4000 × *g*) and separated from spent media. Metabolite extraction was performed by addition of 200 µL cold 2:2:1 acetonitrile:methanol:water. Cell extracts and media were analyzed by HILIC-ESI-qTOF operated in positive mode. See above and SI Appendix for details.

## Data Availability

X-ray data of AgcA_EP4_ are available on the Protein Data Bank with the accession code 9IA1.

## Acknowledgements

This study was supported by a grant from the Natural Sciences and Engineering Research Council of Canada (DG 171359) to LDE. JEM and MZ acknowledges Research England for E3 funding. MEW is the recipient of a Canada Graduate Scholarship-Doctoral. LDE is the recipient of a Canada Research Chair. DJH was funded through NLR under U.S. Department of Energy contract DE-AC36-08GO28308 and the University of Portsmouth. The authors would like to thank Diamond Light Source (Didcot, UK) for beamtime (Proposal Number MX23269) and the staff of beamline I03 for assistance with data collection. We also thank Anne T. Lalande for assistance with molecular biology and protein purification, as well as Dr. Gregg T. Beckham for 4PS.

