## Supplementary material for "Engineering a cytochrome P450 *O*-demethylase for the bioconversion of hardwood lignin"

**This PDF file includes:**

Supporting text
Figures S1 to S18
Tables S1 to S9
SI References

### Supporting Text

#### Protein production and purification

$AgcA_{RHA1}$ and $AgcB_{EP4}$ were produced heterogously as an N-terminal polyHis-tagged proteins as described in (1), with the following modifications. Freshly transformed cells of *E. coli* BL-21 λ (DE3) containing pET28*agcA* or pET28*agcB_EP4* were grown at 37 °C in 1 L Luria broth (LB) containing 50 µg/mL kanamycin sulfate to $OD_{600}$ ~1.0. Following induction with 0.5 mM isopropyl β-D-thiogalactopyranoside, the cells were incubated a further 18 h at 30 °C for AgcA or 20 °C for AgcB. Following homogenization, lysates were chilled on ice and supplemented with protease inhibitors (cOmplete Mini, ½ tablet) and DNAse (~1 mg). N-terminally His-tagged (Ht-) proteins were purified using immobilized metal affinity chromatography (IMAC) (Ni Sepharose 6 fast flow resin, Qiagen). Purification buffers contained 20 mM Tris-Cl, pH 8, 10% glycerol. The protein-loaded resin was washed with an equilibration buffer (10 mM imidazole, 300 mM NaCl), then twice with a wash buffer (50 mM, 300 mM NaCl), and the protein was eluted using 5 mL 250 mM imidazole. Fractions containing tagged protein, as determined by sodium dodecyl sulfate polyacrylamide gel electrophoresis (SDS-PAGE), were pooled. For AgcA, hemin was added dropwise to the pooled fractions to a final concentration of 3:1 hemin:AgcA molar ratio. The protein was dialyzed overnight against 20 mM Tris, pH 8.0, 10% glycerol at 4 °C. Tobacco Etch Virus (TEV) protease was added to approximately 0.1 mg/mL and incubated for 1 h at room temperature with gentle agitation. TEV protease and uncleaved protein were removed by passing the incubation over 1 mL of Ni Sepharose resin. The protein was further purified by passage using a MonoQ 10/100 GL ion-exchange column with an ÄKTA Purifier (GE Healthcare). The protein was eluted using a linear gradient from 0 to 1 M NaCl in 120 mL of 20 mM Tris, pH 8.0. Fractions with a $R_z$ ($Abs_{418\ nm}/Abs_{280\ nm}$) > 1 were pooled and exchanged into 20 mM Tris, pH 8.0, 10% glycerol, and flash frozen dropwise as ~10 µL beads in liquid $N_2$.

For AgcB, all steps following lysis were conducted anaerobically inside a Labmaster Model 100 glovebox (Mbraun). IMAC was conducted as described above for AgcA. Yellow-brown colored fractions were pooled and exchanged into 20 mM Tris, pH 8.0, 10% glycerol using a 10 mL Amicon Stirred cell fitted with a 30 kDA MWCO Ultracel Ultrafiltation disc and flash frozen as ~10 µL drops in liquid $N_2$.

For crystallography, $AgcA_{EP4}$ was produced using *E. coli* Lemo21 (DE3) and Terrific broth (TB) media. At an $OD_{600}$ of 0.6, 0 IPTG and 5-aminolevulinic acid were added to 5 mM and 100 mg $L^{-1}$, respectively. The cultures were incubated overnight at 17 °C. Cell pellets were harvested by centrifuging at 5,000 × *g* for 20 min. Pellets were either frozen at -20 °C for storage or taken onto purification. Cells were lysed using sonication in 20 mM HEPES, 200 mM NaCl, pH 8.0 and then centrifuged at 50,000 × *g* for 1 h to pellet cell debris. Subsequent purification steps were as previously described (2). Purified protein was then either used immediately or flash frozen in liquid $N_2$ and stored at -80 °C for later use.

For all other studies, $AgcA_{EP4}$ and its variants were prepared as previously described (2). Briefly, Ht-$AgcA_{EP4}$ was produced heterologously in *E. coli* BL-21 λ (DE3). Protein production was initiated by adding IPTG to 1-L of cultured *E. coli* bearing the appropriate expression plasmid (**Table S9**). Cells were harvested by centrifugation and lysed at 4 °C using an EmulsiFlex-C5 homogenizer (Avestin). The tagged protein was purified from the cell extract using IMAC and size-exclusion chromatography. AgcA and derivatives were reconstituted with heme by dropwise addition of 50 mM hemin (to three molar equivalents) in 0.1 M NaOH while stirring. The His tag was removed by incubation with TEV protease (50:1 molar ratio). The mixture was dialyzed overnight against 20 mM Tris, pH 8.0, 10 % glycerol at 4 °C. The cleaved and uncleaved Ht-AgcA were separated by passing the dialyzed mixture over the Ni resin. AgcA and derivatives were further purified using a MonoQ 10/100 GL column and an ÄKTA Purifier (GE Healthcare). The protein was concentrated to approximately 10 mg $mL^{-1}$, flash-frozen as ~10 µL drops, and stored at -80 °C.

#### *Protein analysis*

AgcA heme concentration was calculated by a carbon monoxide (CO) binding assay as described previously (3). Briefly, CO binding was determined by reducing AgcA (diluted in 20 mM MOPS, $I$ = 0.1 M, pH 7.2) using a few grains of sodium dithionate. CO was bubbled into the solution of reduced enzyme for approximately 60 s at a rate of ~one bubble per second. AgcB FAD concentration was determined by dilution of AgcB in 20 mM MOPS, I = 0.1 M, pH 7.2, boiling for 10 min in the dark, then calculating FAD concentration using absorbance at 450 nm and ε = 11.3 $mM^{-1}$ $cm^{-1}$. For all further enzyme activity assays, [AgcA] and [AgcB] were defined by [heme] and [FAD], respectively.

#### *Crystallography*

Crystals in complex with 20 mM 4-ethylguaiacol (4EG) were grown at 20 °C in 20% PEG 6000, 0.2 M $CaCl_2$ and 0.1 M MES, pH 6.0 with a protein concentration of 10 mg $mL^{-1}$, with multiple rounds of seeding required to produce crystals that diffracted to 1.82 Å. Diffraction data were collected at beamline I03 at the Diamond Light Source (Didcot, UK). The unmerged data file from Dials (4) at Diamond were reduced using to Aimless (5) to 1.82 Å and the structure was solved by molecular replacement with Phaser (6) using PDB: 5NCB as the search model, and refined using Refmac5 (7). The structure was manually refined using Coot (8) and validated with MolProbity (9). Diffraction data statistics are shown in **Table S1**. Structural figures were generated in PyMOL (Schrödinger, LLC). RMSD values were calculated using the combinatorial extension (CE) alignment plugin in PyMOL (10). The structure of $AgcA_{EP4}$ in complex with 4-EG are available at the Protein Data Bank (PDB) under the accession code: **9IA1**. The structure of $AgcA_{RHA1}$ was predicted with the Swiss-Model homology-modelling server with the structure of $AgcA_{EP4}$ (9IA1) as a template (11).

#### *Determination of binding constants ($K_D$) and monoxy-ferrous heme state*

UV-Vis spectroscopy was conducted with a Cary 60 spectrophotometer equipped with a thermostatted cuvette holder. Scans of purified proteins diluted in 10 mM MOPS, *I* = 25 mM, pH 7.2 at 25 °C were taken between 250 and 700 nm. This buffer was prepared by titrating 5 mN NaOH with MOPS to pH 7.2 (~10 mM MOPS, *I* = 5 mM). Ionic strength was increased by adding the appropriate amount of NaCl. Heme concentration was calculated as described previously (3). Dissociation constants ($K_d$) were determined by titration of ligands diluted into enzyme buffer from 1 M stocks prepared in DMSO into AgcA diluted in 10 mM MOPS, *I* = 25 mM, pH 7.2. Spectra were recorded after addition of substrate and subtracted from the spectrum of the resting-state enzyme. The absorbance difference between the peak and trough of the subtracted spectra was plotted against ligand concentration and $K_d$ values were determined by fitting a hyperbolic equation to the data as described previously (12, 13). In the case of 4PG binding by WT AgcA, the data were fit to the quadratic tight-binding equation (3). The proportion of high-spin heme iron was determined using the approximate peak-to-trough extinction coefficient (ε ≈ 100 $mM^{-1}cm^{-1}$) (13). The monoxy-ferrous species of the heme iron was formed by reducing a mixture of 2 µM of AgcA with 100 µM 4EG in CO-saturated buffer with either 5 µM AgcB and 350 µM NADH or a few grains of sodium dithionite. Buffer was saturated with CO by gently bubbling for 20 min.

#### *Coupling assays*

Activity assays for coupling were conducted using 1 µM of AgcA and AgcB in 200 µL 10 mM MOPS, pH 7.2 (*I* = 25 mM), 25 °C. Ionic strength was adjusted using NaCl as described above. The reaction was initiated by adding NADH to 350 µM. For HPLC analysis, enzymatic reactions were acidified by adding acetic acid to 10% final concentration, centrifuged (16,000 × *g* for 10 min), and filtered through a 0.2 µm PTFE membrane. Samples were run over a Luna 5 µm C18(2) 100 Å 150 × 3 mm column (Phenomenex) at 0.7 mL $min^{-1}$ by a Waters 2695 separation HPLC module. Samples were eluted with a 16.8 mL linear gradient from 1% methanol plus 0.1% formic acid in water to 100% methanol plus 0.1% formic acid and monitored at 280 nm with a Waters 2996 photodiode array detector. Concentrations were determined by interpolation on a standard curve of 0 to 1 mM of authentic standard. NADH concentration was measured by absorbance at 340 nm, ε = 6.22 $mM^{-1}$ $cm^{-1}$. Coupling was calculated from the ratio of rate of aromatic turnover, as determined by HPLC, and the rate of NADH oxidation, determined spectroscopically. Formaldehyde was determined *via* a trypophan-based spectrophotometric assays as described previously (3). Briefly 200 µL of 0.1% tryptophan (in 50% ethanol), 200 µL 90% sulfuric acid and 40 µL of 1% $FeCl_3$ were added to 200 µL of quenched enzymatic reaction in wells of a 24-well plate. The solution was incubated for 60 min at 70 °C on a heat block. Concentration of formaldehyde was calculated by absorbance at 575 nm and interpolation of a standard curve of 0 – 1330 mM of formaldehyde.

#### *Apparent steady-state kinetic parameters*

For $AgcA_{EP4,}$ the rate of 4PG and guaiacol turnover were determined using a coupled assay with AphC or XylE, as described previously (2). For $AgcA_{RHA1}$, NADH depletion rate was measured spectroscopically (ε = 6.22 $mM^{-1}$ $cm^{-1}$) and initial rate was adjusted by coupling to yield the estimated aromatic turnover rate. Apparent steady-state kinetic parameters were calculated by fitting Michaelis-Menten equations to initial velocity of reactions at various concentrations of aromatic substrate (LEONORA).

For detection of reaction products, quenched reaction mixtures were analyzed by liquid chromatography-mass spectrometry (LC-MS) using an Agilent 1290 Infinity II UHPLC in line with an Agilent 6546 Q-TOF equipped with a dual AJS ESI source operating in positive and negative ionization modes. A sample (2 µL) was injected onto a Zorbax Eclipse Plus C18 Rapid Resolution HD, 2.1 × 50 mm × 1.8 µm column and run on a 12 min linear gradient from 5% to 100% solvent B at 0.25 mL $min^{-1}$. Solvent A was 0.1% formic acid in water, and solvent B was 0.1% formic acid in acetonitrile (ACN). MS parameters were as follows: capillary voltage, 3500 V; nozzle voltage, 500 V; drying gas temp, 300°C; drying gas flow rate, 10 L $min^{-1}$; sheath gas temperature, 350°C; sheath gas flow rate, 12 L $min^{-1}$; nebulizer pressure, 45 psi; fragmentor voltage, 100 V. The MS parameters used for negative mode were the same as positive with the following differences: nozzle voltage, 1000 V; drying gas temp, 250 °C; sheath gas temperature, 300°C; nebulizer pressure, 40 psi. MS/MS was collected on selected ions with 10, 20, and 40 V collision energies. Data were collected and analyzed using MassHunter Workstation version 10.

For detection of pathway metabolites, cell extracts and spent media were analyzed LC-MS on the same system. Samples (2 µL) were injected onto an InfinityLab Poroshell 120 HILIC-Z column (100 × 2.1 × 2.7 µm) and resolved using a 10 min linear gradient from 90 to 60% solvent B at 0.25 mL/min. Solvent A was 10 mM ammonium acetate, pH 9 and solvent B was 10 mM ammonium acetate, pH 9 in 90% acetonitrile. MS parameters were as follows: capillary voltage, 3500 V; nozzle voltage, 500 V; drying gas temp, 250 °C; drying gas flow rate, 10 L/min; sheath gas temperature, 300 °C; sheath gas flow rate, 12 L/min; nebulizer pressure, 40 psi; fragmentor voltage, 100 V. Data were processed using Agilent MassHunter Profinder and Agilent TOF Quantitative Analysis Version 10.

#### *Growth assays*

Rhodococcus was grown at 30 °C. For the growth experiment, single colonies of RHAMW30, RHAMW31 and RHAMW32 were cultured in 2 mL of LB, supplemented with 0.5 mM 4EG. When the culture attained OD ~0.5, the cells were harvested by centrifugation (3000 × *g*), washed twice with M9 minimal media and suspended in 500 µL M9 minimal media supplemented with goodies (M9G) and the appropriate substrate. Aromatic substrates were prepared and used as 1 M stocks in dimethyl sulfoxide. Cells were incubated at 30 °C for 42 h in 48-well plates shaking at 200 rpm. OD600 was measured using a TECAN Spark.

#### *Meta-cleavage activity of lysates*

Colonies of RHAMW32 were cultured in 2 mL LB until stationary phase (OD600 ~ 15). These cells were used to inoculate 50 mL 10 mM glucose in M9G and incubated overnight. At OD600 ~1.5, cells were harvested by centrifugation (3000 × *g*) and suspended to OD600 0.25 in 1 mL 10 mM glucose in M9G supplemented with either 1 mM 4PG, 1 mM 4PS or no additional substrate in 24-well plates. Cells were incubated for 20 h with shaking at 200 rpm, then harvested by centrifugation (13 000 × *g*), and suspended in 500 µL of lysis buffer (100 mM $NaPO_4$, pH 8, 10% glycerol). The suspension was added to 2 mL freezer tubes with ~100 µL 0.1 mm Zr/Si beads and ~50 µL of 0.5 mm glass beads. Cells were lysed using a FastPrep-24™ Classic Bead Beating Grinder (MP Biomedicals™) and three 45 s cycles at 6 m/s, then clarified by centrifugation (13 000 × *g*).

Activity was measured spectrophotometrically using a Cary 60 spectrophotometer equipped with a thermostatted cuvette holder. Aliquots of the clarified lysates (200 µL) were added to a quartz cuvette and equilibrated to 25 °C. The reaction was initiated by adding 4-methylcatechol (4MC) to 200 µM and followed for 10 min. The rate of reaction was calculated using an extinction coefficient at 382 nm of 29 $mM^{-1}$ $cm^{-1}$ for 5-methyl HODA (14). Rates were normalized to total protein in the lysates, measured using a Pierce™ BCA Protein Assay Kit (Thermo Scientific™).

#### *Resting cell assays*

Single colonies of RHAMW30, RHAMW31 and RHAMW32 were cultured in 2 mL LB. These were grown overnight, then used to inoculate 100 mL LB. When cultures reached OD600 ~3, cells were harvested by centrifugation, washed twice with M9, and suspended in 500 µL 0.2% glucose in M9G containing either 0.2 mM 4PG or 0.3 mM 4PS at an OD600 of 10 in a 48 well plate. Media was sampled (30 µL) at 30, 60 and 120 min, and remaining substrate was quantified by HPLC using the method described above. To detect metabolites of the Agc/Aph pathway in RHAMW32, a single colony of RHAMW32 was cultured in 2 mL LB. This culture was used to inoculate 50 mL 10 mM glucose in M9G. Cells were collected at an OD600 of 0.5-2, washed twice using M9, and suspended in M9 to OD600 1. Fifteen mL aliquots of the concentrated culture were incubated with 1 mM 4EG for 4 h at 30 °C, collected by centrifugation, and suspended in 5 mL 10 mM glucose M9G supplemented with 1 mM 4PG or 4PS to an OD600 of ~2. Resting cell suspensions were incubated with 0.5 mM 4PS or 4PG and shaking at 30 °C for 6 h, then harvested by centrifugation (4000 × *g*) and separated from spent media. Metabolites were extracted from cells by addition of 200 µL cold acetonitrile:methanol:water (2:2:1). Cell extracts and media were analyzed by HILIC-ESI-qTOF operated in positive mode as described above.

```
AgcA_EP4     ----MTSTHS-----FIDEITIEELEADPYPFYERLRKEAPIAYVPALGMYIVSTKELCAEI---SKDD
AgcA_RHA1    ----MTTSTT-----WIESITMEELDRDPNPIYDRLRREAPVAFVPAVGMHVVASRDLCLQIAQDSETW
GcoA         MTTTERPDLA-----WLDEVTMTQLERNPYEVYERLRAEAPLAFVPVLGSYVASTAEVCREVAT-SPDF
SyoA         --MTTKHTTAGDTQEWLATVTVEQLENDPYPIFERLRREAPVAWIPAAHAWVASTWEACRTIADDATNF


AgcA_EP4     ANWPAVISAAGGRTFGPQALLNTNGDEHRNLRDMVEPHLQPSAVDKYIDDLVRPFARQRIAEFENDGHA
AgcA_RHA1    ---STVIAPSGGRTFGKGTVLAANGEQHEKIREWIDPQLRPSAVDSYVEALVRPQARSLLEGIEDLGAA
GcoA         ---EAVITPAGGRTFGHPAIIGVNGDIHADLRSMVEPALQPAEVDRWIDDLVRPIARRYLERFENDGHA
SyoA         ---RGGTSPMHERVLGTDHILGAEGETHQDLRAAVDPPLKPRAFRPLLEEQVRPTVRRYLEAIRGQGKA


AgcA_EP4     DIVAAYCEPVSVRALGDLLGLGDVSTEKLREWFHNLSVSFTNAAVDENGEFANPEGFAPGDRAKAEIIA
AgcA_RHA1    DIQEAYFAPISVRSVGDLMGLTEIPSETLVRWFETLAQSYGNAEVDENGNFANPGPFEAGDRVKAEIVA
GcoA         ELVAQYCEPVSVRSLGDLLGLQEVDSDKLREWFAKLNRSFTNAAVDENGEFANPEGFAEGDQAKAEIRA
SyoA         ELMADYFEPISVRCVGDVIGLTDVDSDTLRRWFHALARGIANTAMDAEGRFTNPGGFAPADEAGAEIRE


AgcA_EP4     HVDPKIDKWIVEPDHSAISHWLHDGMPEGQTRSRDVIYPNLYVFLLGAMQEPGHAMATTLAGLFSRPDQ
AgcA_RHA1    AVGPMLDHWTEHPDHTLISHWLHDGMPDGQVRDRSEIYPNIYVFLLGALQEPGHVMTTTLAGLFQHPDQ
GcoA         VVDPLIDKWIEHPDDSAISHWLHDGMPPGQTRDREYIYPTIYVYLLGAMQEPGHGMASTLVGLFSRPEQ
SyoA         VLEPLVAKLSAEPDGSALSHYLHHGRPHGDPRTLEQLLPSLKVIILGGLQEPGHQCAATFLGLTTRPEQ


AgcA_EP4     LERVIDDPTLIPRAASEGMRWVAPIWSAAVKRAAREVTVGGVTLPEGSIVMLSYGSANQDENAYNAPTE
AgcA_RHA1    LERVIDDPTLIPRAVNEGARWVAPIWSAAVKVAGRDVTIGGIDLPTGTPVMLAYGSANRDESVWENAEA
GcoA         LEEVVDDPTLIPRAIAEGLRWTSPIWSATARISTKPVTIAGVDLPAGTPVMLSYGSANHDTGKYEAPSQ
SyoA         LKRVTEDATLLPRALTEGLRWMSPVFSASSRLPLREITMGEATMRPGQTVWLSYGSANRDEAVFDRPDV


AgcA_EP4     YDLDRALVPNMTFGGGKHACAGTYFANAVVRIGLEELLEAIPNIERDETHE-VDFWGWGFRGPKQLFVK
AgcA_RHA1    YEIDRPIMPHLAFGAGNHACAGTYLGTAIVRIALEALFETIPNIEPD-PDRAPQFWGWTFRGPQGLHVT
GcoA         YDLHRPPLPHLAFGAGNHACAGIYFANHVMRIALEELFEAIPNLERD-TREGVEFWGWGFRGPTSLHVT
SyoA         FDLDRATHPHLAFGTGRHLCSGSAYAPQVARIALEELFTAFPSIRLD-PAHEVPVWGWLFRGPQRLDVL


AgcA_EP4     WEV----
AgcA_RHA1    WEV----
GcoA         WEV----
SyoA         WDGGGSG
```

**Figure S1: Alignment of select members of the CYP255A subfamily**. Selected sequences are AgcA (2), $GcoA_{Amy}$ (3), and SyoA (15). Residues are highlighted according to their potential roles in the following functions: yellow, specificity for the alkyl side chain; green, specificity for syringol. Sequences were aligned using Clustal Omega (16).

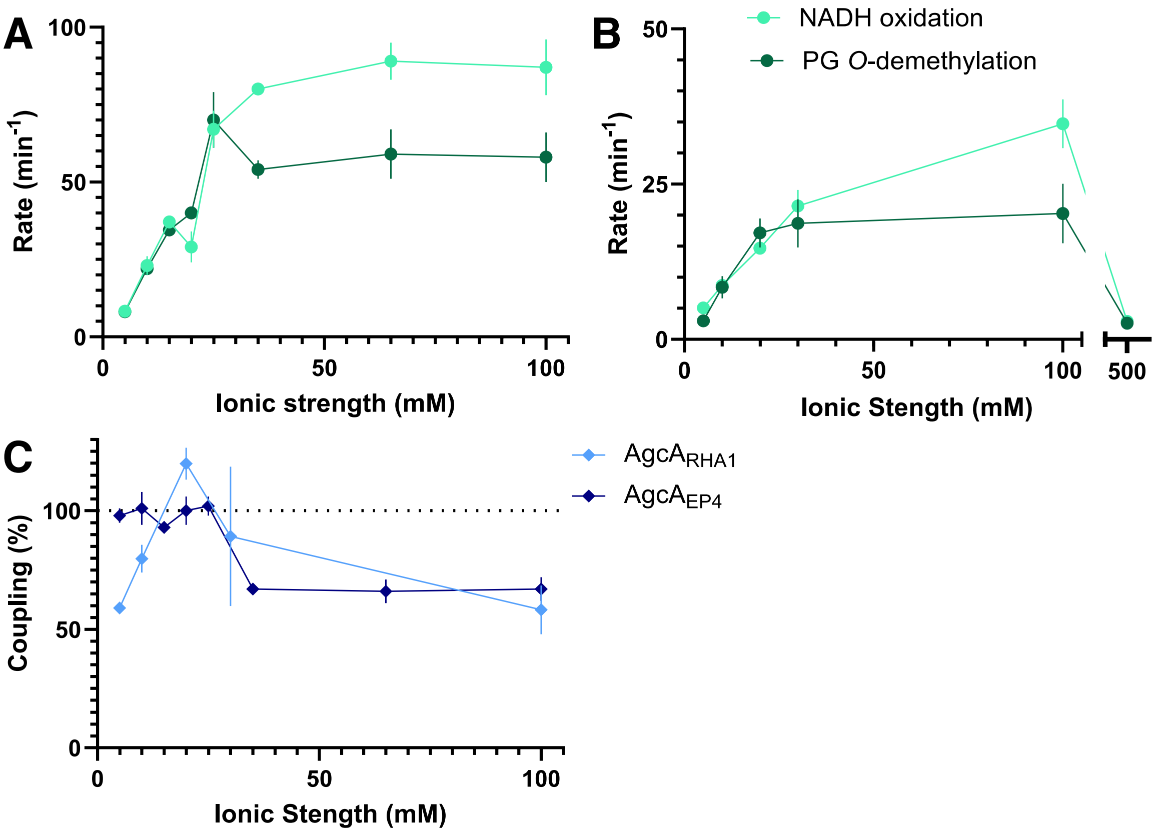


**Figure S2: Influence of ionic strength on AgcA activity and coupling.** Experiments were performed using **(A)** 1 µM AgcA$_{EP4}$ or **(B)** 1 µM AgcA$_{RHA1}$ and 1 µM AgcB, 100 µM 4PG and 350 µM NADH in 10 mM MOPS, pH 7.2 (adjusted to the indicated ionic strength with NaCl) at 25 °C. Appearance of the *O*-demethylated product was measured by HPLC and oxidation of NADH was measured by spectroscopy. (**C**) Coupling efficiency (calculated as (rate of catechol appearance)/(rate of NADH depletion)) of the AgcA$_{RHA1}$ (light blue) and AgcA$_{EP4}$ (dark blue) systems.

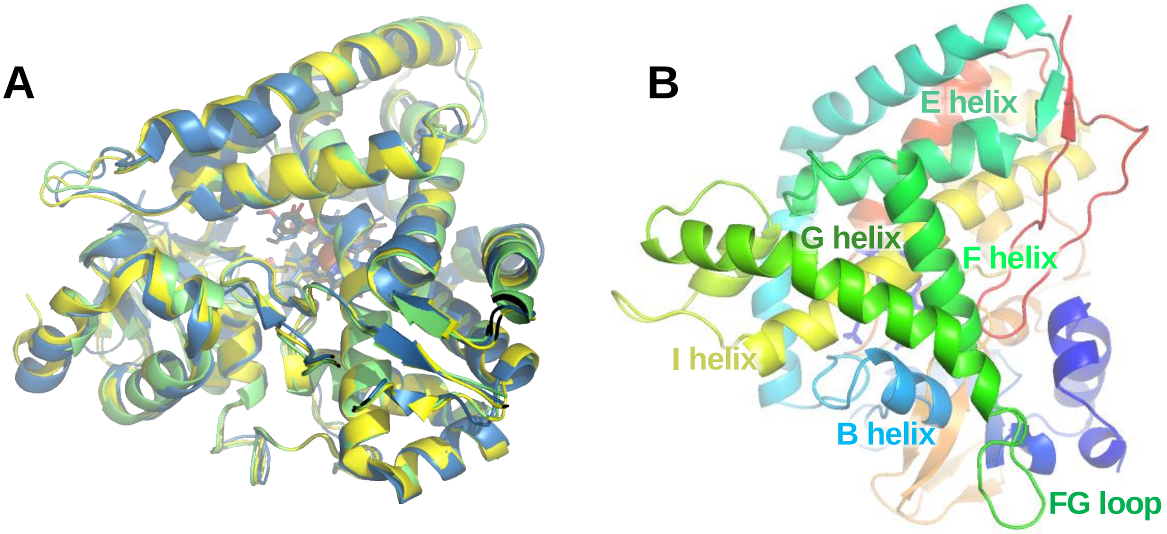


**Figure S3: Overall properties of the AgcA structure.** (**A**) AgcA$_{EP4}$ (green, 9IA1), GcoA (yellow, 5NCB) and SyoA (blue, 8U19) in their substrate-bound (closed) forms share the canonical fold of P450s, including (**B**) helices which make up the substrate-binding pocket.

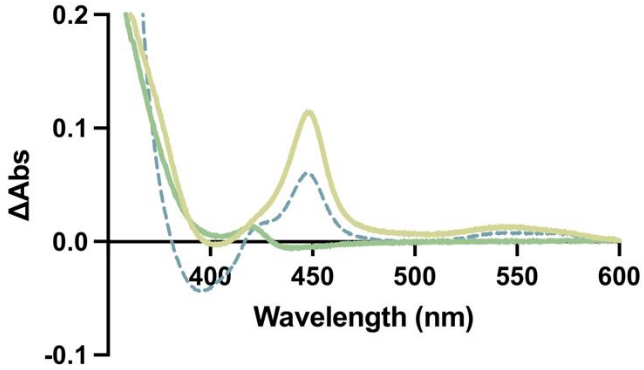


**Figure S4: Formation of the ferrous-CO complex in AgcA$_{EP4}$ F166N variant.** Difference spectra of 2 µM AgcA$_{EP40}$ WT (light green) and F166N (dark green)) ± 350 µM NADH when combined with 100 µM 4PG and 5 µM AgcB in CO-saturated 10 mM MOPS, pH 7.2, *I* = 25 mM at 25 °C. The CO-difference spectrum of AgcA F166N was also recorded under similar conditions using sodium dithionite instead of NADH and AgcB (dashed blue).

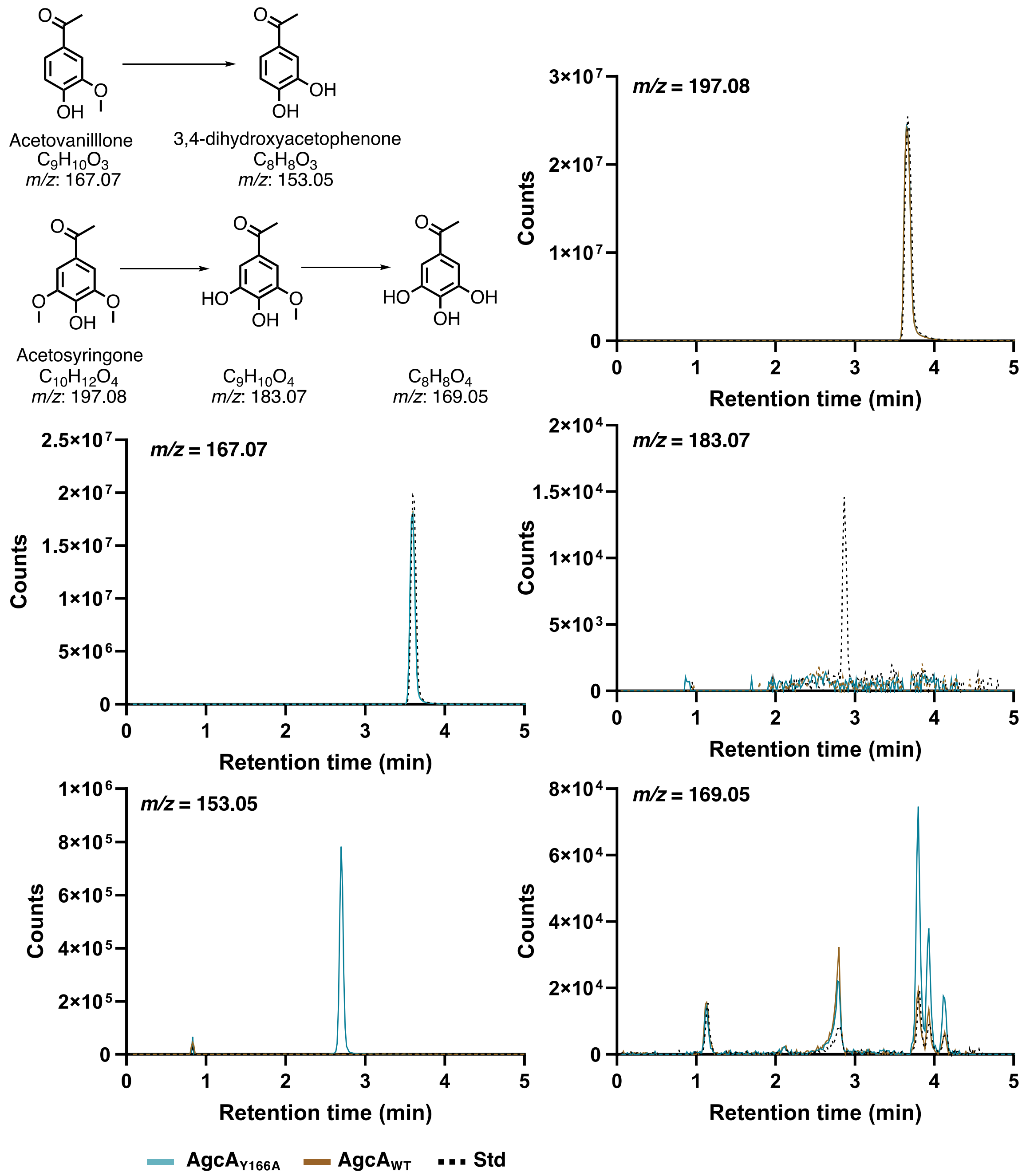


**Figure S5: *O*-Demethylation of acetophenones by AgcA$_{RHA1}$.** Acetovanillone (100 µM) or acetosyringone (200 µM) were incubated with 1 µM AgcAB WT (blue EIC) and Y166A (brown EIC), 350 µM NADH and 1000 U/mL catalase for 20 h at 25 °C. EICs shown are of predicted *m*/*z* values in positive mode of acetophenones and their *O*-demethylation products.

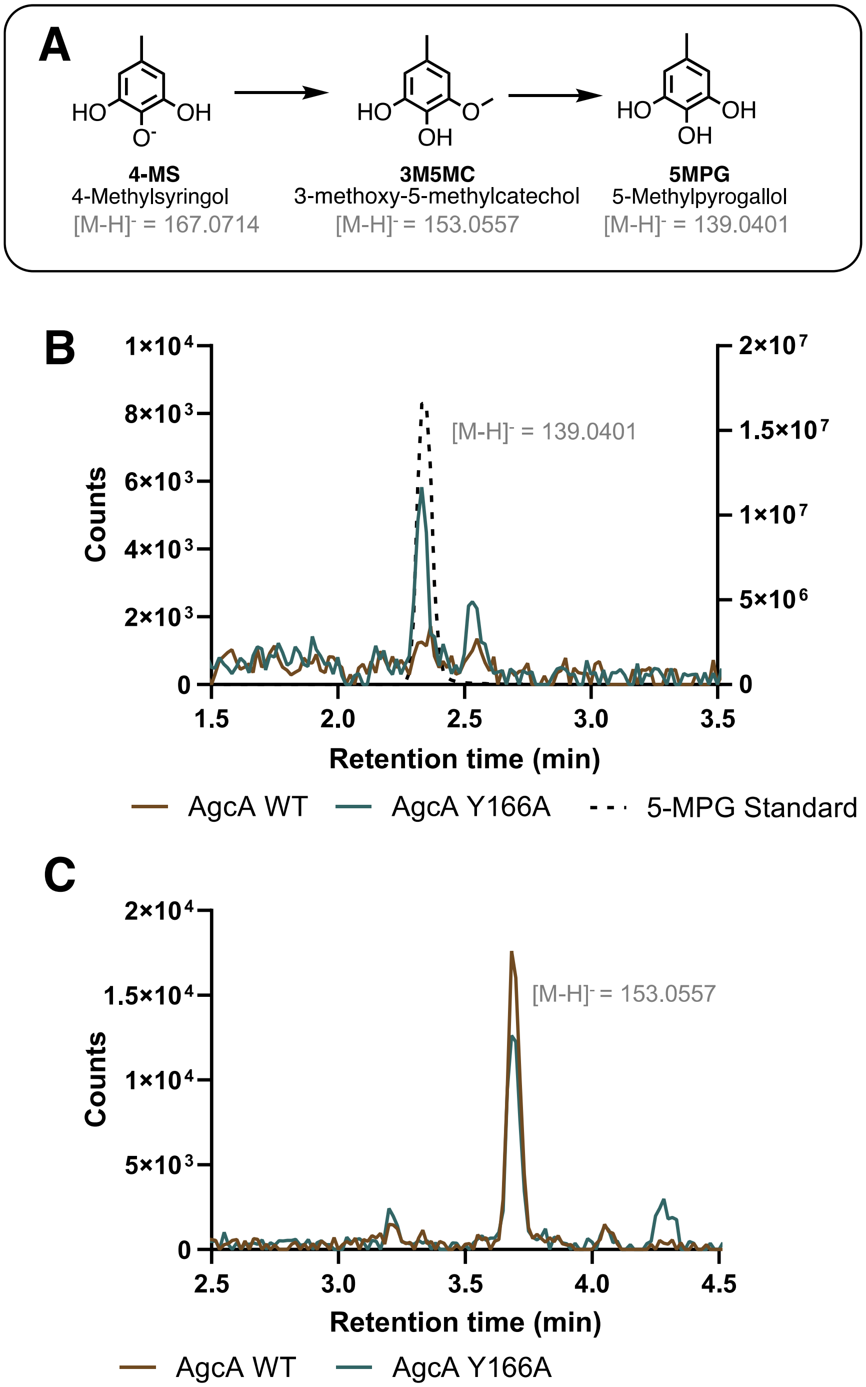


**Figure S6: *O*-Demethylation of 4MS by AgcA$_{RHA1}$ WT and Y166A: (A)** Predicted *O*-demethylation scheme of 4MS, with products and *m/z* of the deprotonated monoisotopic mass. Reactions containing 1 µM WT or Y166A AgcA$_{RHA1}$, 1 µM AgcB, 100 µM 4MS and 350 µM NADH in 10 mM MOPS, pH 7.2, *I* = 25 mM were incubated for 10 min at 25 °C then quenched. The quenched reaction was analyzed by LC-MS-ESI (-). Extracted ion chromatograms (EICs) of *m/z* corresponding to the demethylation products 5MPG **(B),** 3M5MC **(C)** and for the WT (brown) and Y166A mutant (teal), alongside 100 µM of the authentic 5MPG (dashed black). For (**B**), the EICs from the reaction mixtures are plotted on the left axis and the standard on the right axis

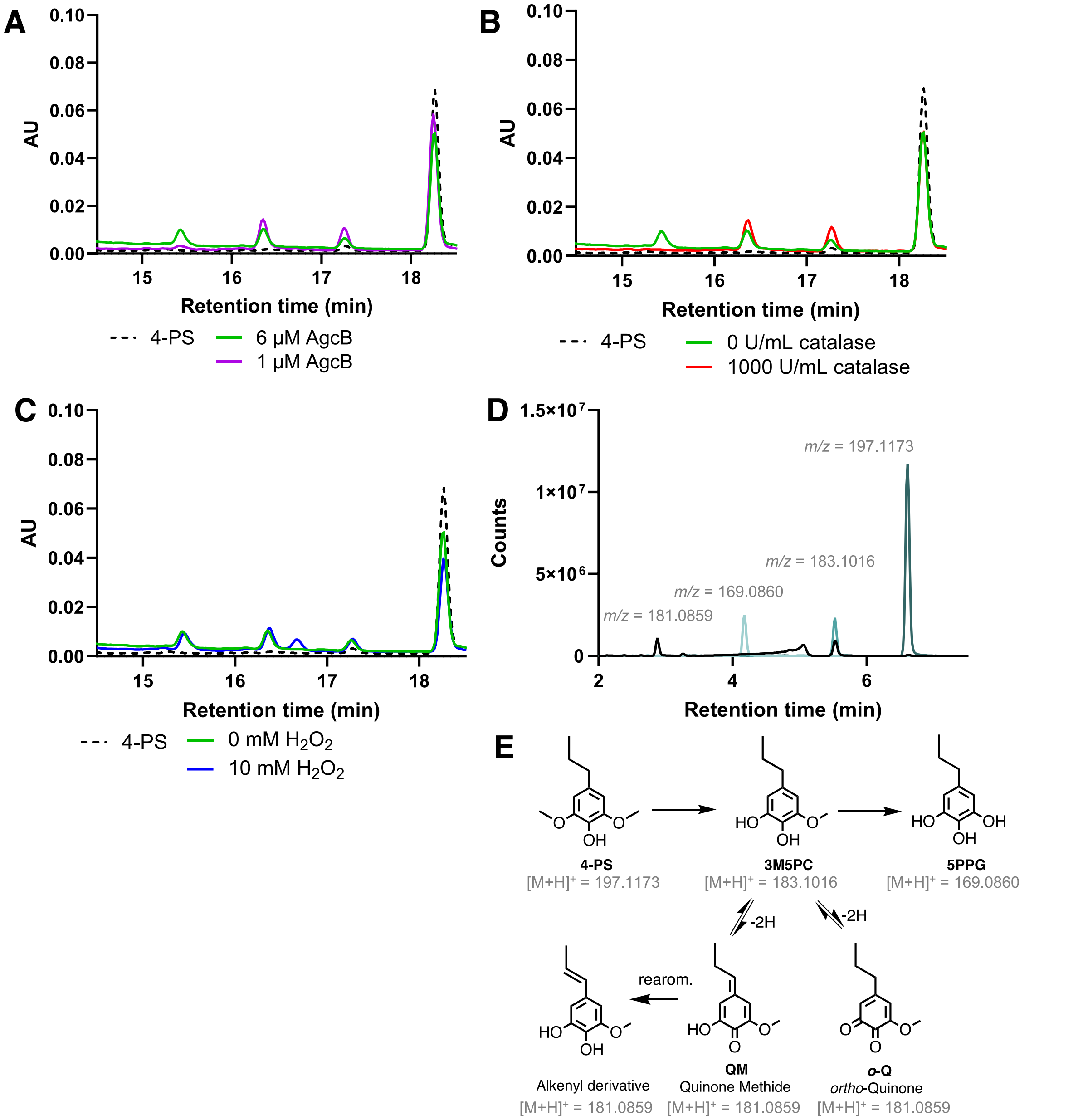


**Figure S7: Effect of reactive oxygen species on the *O*-demethylation of 4PS by AgcA$_{RHA1}$ Y166A.** Reactions containing 1 µM AgcA Y166A, 6 µM AgcB, 100 µM 4PS, and 350 µM NADH in 10 mM MOPS, pH 7.2, *I* = 25 mM (green trace) and were quenched after 10 min incubation at 25 °C with the following modifications: **(A)** reduction to 1 µM AgcB (purple); **(B)** addition of 1000 U/mL catalase (red); and **(C)** addition of 10 mM $H_2O_2$ (blue). **(D)** LC-MS-ESI (+) analysis of the quenched reaction. Extracted ion chromatograms (EICs) of *m/z* corresponding to the substrate 4PS (dark blue) and predicted *O*-demethylation products, 3M5PC (aqua blue) and 5PPG (light blue), and the oxidized catechol, black. (E) proposed reaction scheme for the generation of quinones and stable isomers from 3M5PC, with predicted *m/z* of the protonated monoisotopic mass.

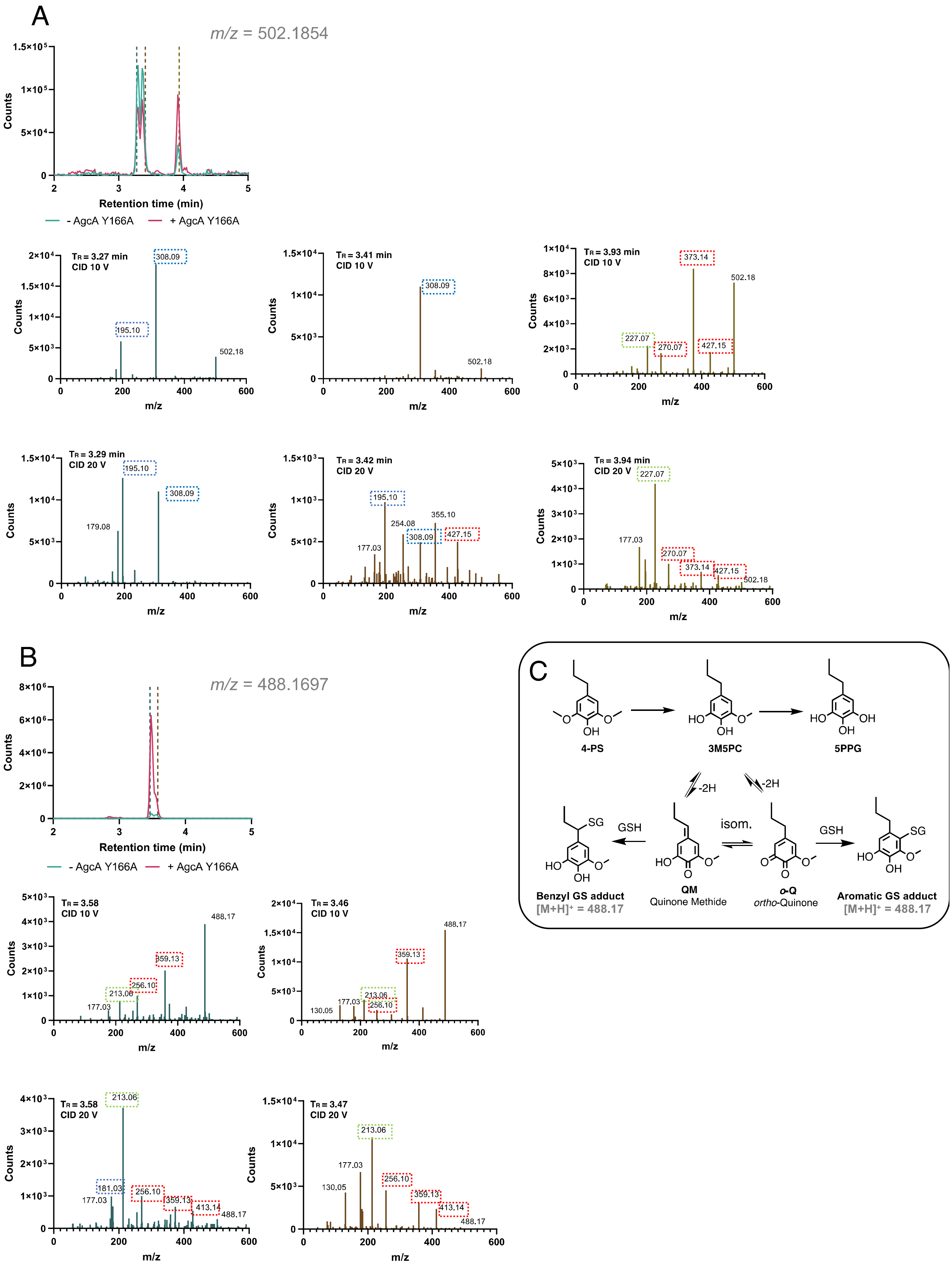
A
m/z = 502.1854
Counts
Retention time (min)
- AgcA Y166A
+ AgcA Y166A
TR = 3.27 min
CID 10 V
TR = 3.41 min
CID 10 V
TR = 3.93 min
CID 10 V
TR = 3.29 min
CID 20 V
TR = 3.42 min
CID 20 V
TR = 3.94 min
CID 20 V
m/z
B
m/z = 488.1697
TR = 3.58
CID 10 V
TR = 3.46
CID 10 V
TR = 3.58
CID 20 V
TR = 3.47
CID 20 V
C
4-PS
3M5PC
5PPG
-2H
GSH
isom.
Benzyl GS adduct
[M+H]+ = 488.17
QM
Quinone Methide
o-Q
ortho-Quinone
Aromatic GS adduct
[M+H]+ = 488.17

**Figure S8: Formation and analysis of GS adducts**. Reactions containing 1 µM AgcA Y166A, 6 µM AgcB, 10 mM $H_2O_2$, 100 µM 4PS and 350 µM NADH in 10 mM MOPS, pH 7.2, *I* = 25 mM were incubated for 10 min at 25 °C in the presence of 20 mM glutathione. LC-MS-ESI (+) analysis of the quenched reaction. (EICs) of *m/z* corresponding to the **(A)** substrate-GS adduct and **(B)** catechol-GS adduct for the full reaction (pink) and no-enzyme (AgcAB) control (teal). Also shown are MS/MS analysis of the parent ion *m/z* = 488.1697 **(A)** or 502.1854 **(B)** for each peak. Characteristic fragments (17) of the GS moiety are highlighted in red, and predicted fragments from the benzyl core are highlighted in blue. Fragments which form from the benzylic adduct are highlighted in blue and of the benzylic or aromatic adduct are in green (18). **(C)** Proposed reaction scheme for formation of the GS adduct of oxidized catechol quinones with predicted *m/z* of the protonated monoisotopic mass of the GS adduct.

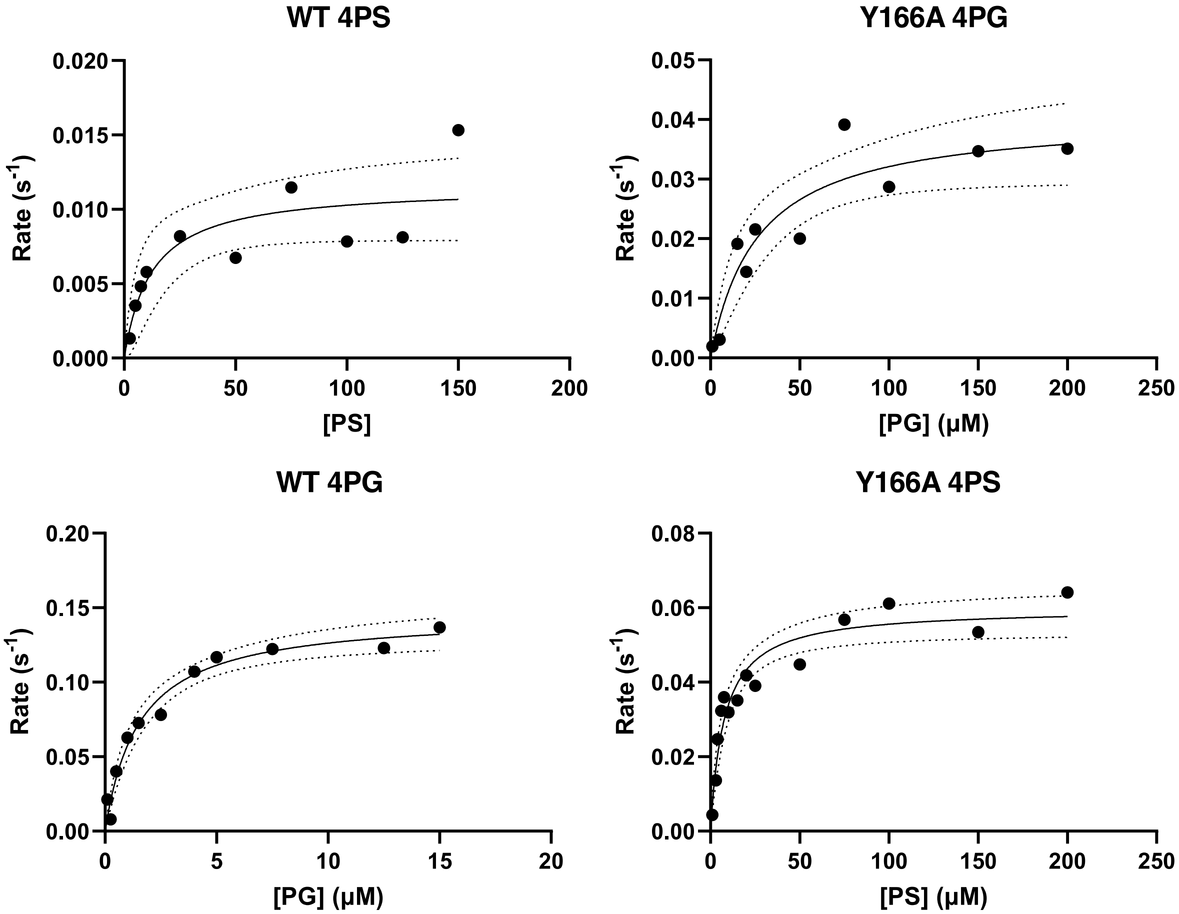


**Figure S9: Michaelis-Menten curves of $\text{AgcA}_{\text{RHA1}}$ WT and Y166A**. Reactions contained 0.2-1 µM AgcAB, 1000 U/mL catalase, 175 µM NADH and 0-250 µM substrate in 10 mM MOPS, pH 7.2, *I* = 25 mM. Reaction rate was measured over a 20 s window in the first 1-3 mins of incubation at 30 °C. For WT 4PG, WT Y166A 4PS, reaction rates were determined spectrophotometrically by measuring NADH oxidation and corrected for coupling (45% in the case of WT at [4PS] > 10 µM). For all other samples, substrate depletion (for 4PS) or product appearance (for 4PC) were measured by LC-MS. Curves represent best fit of the Michaelis-Menten equation to the data (solid line).

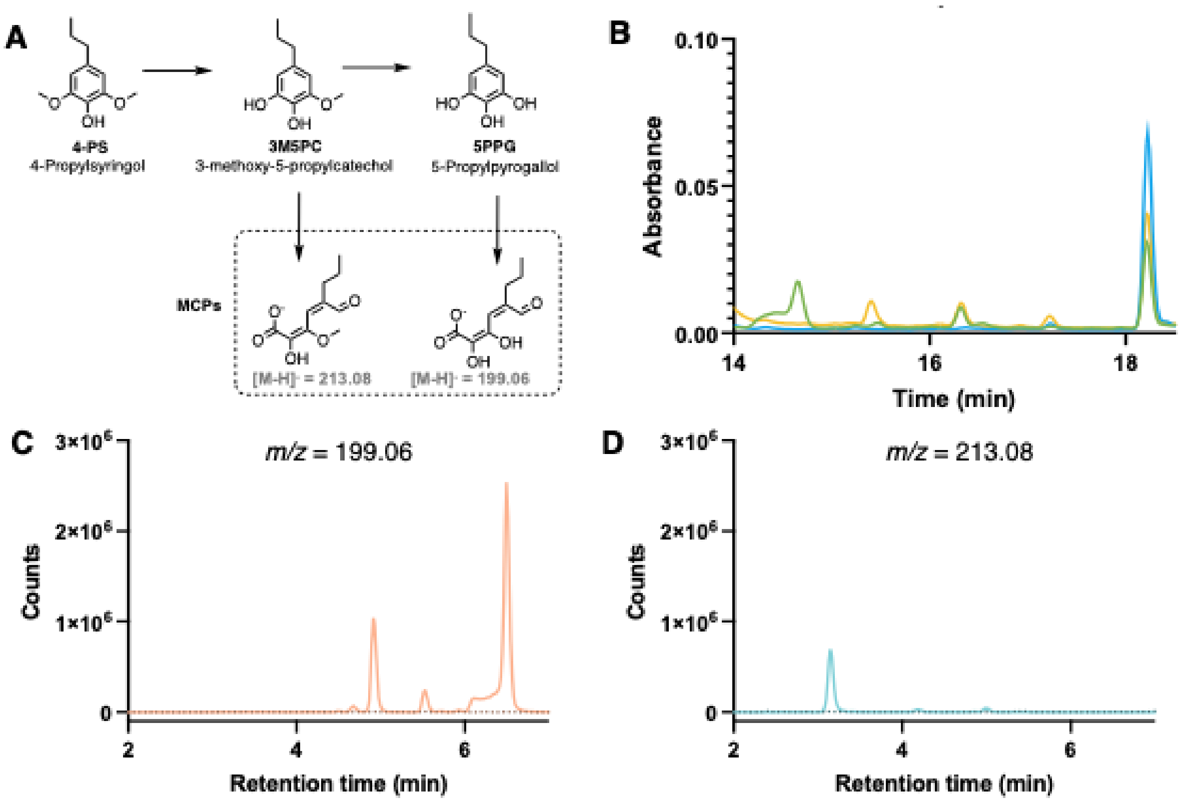


**Figure S10: Cleavage of 4PS *O*-demethylation products by AphC. (A)** The predicted *O*-demethylation products of 4PS and their *meta*-cleavage by AphC. The indicated *m/z* values are those calculated for the deprotonated monoisotopic MCPs. Reactions containing 1 µM AgcA Y166A, 1 µM AgcB, 100 µM 4PS, 15 µM AphC and 350 µM NADH in 10 mM MOPS, pH 7.2, *I* = 25 mM were incubated at 25 °C for 30 min then quenched. **(B)** HPLC chromatogram of the full reaction (green), 10 min incubation excluding AphC (yellow), and 10 min incubation excluding AgcAB (blue). **(C)** LC-MS-ESI (-). Extracted ion chromatograms (EICs) of *m/z* corresponding to the MCPs of 3M5PC **(C)** and 5PPG **(D)** for the +AphC (solid lines) and -AphC (dotted lines) reactions.

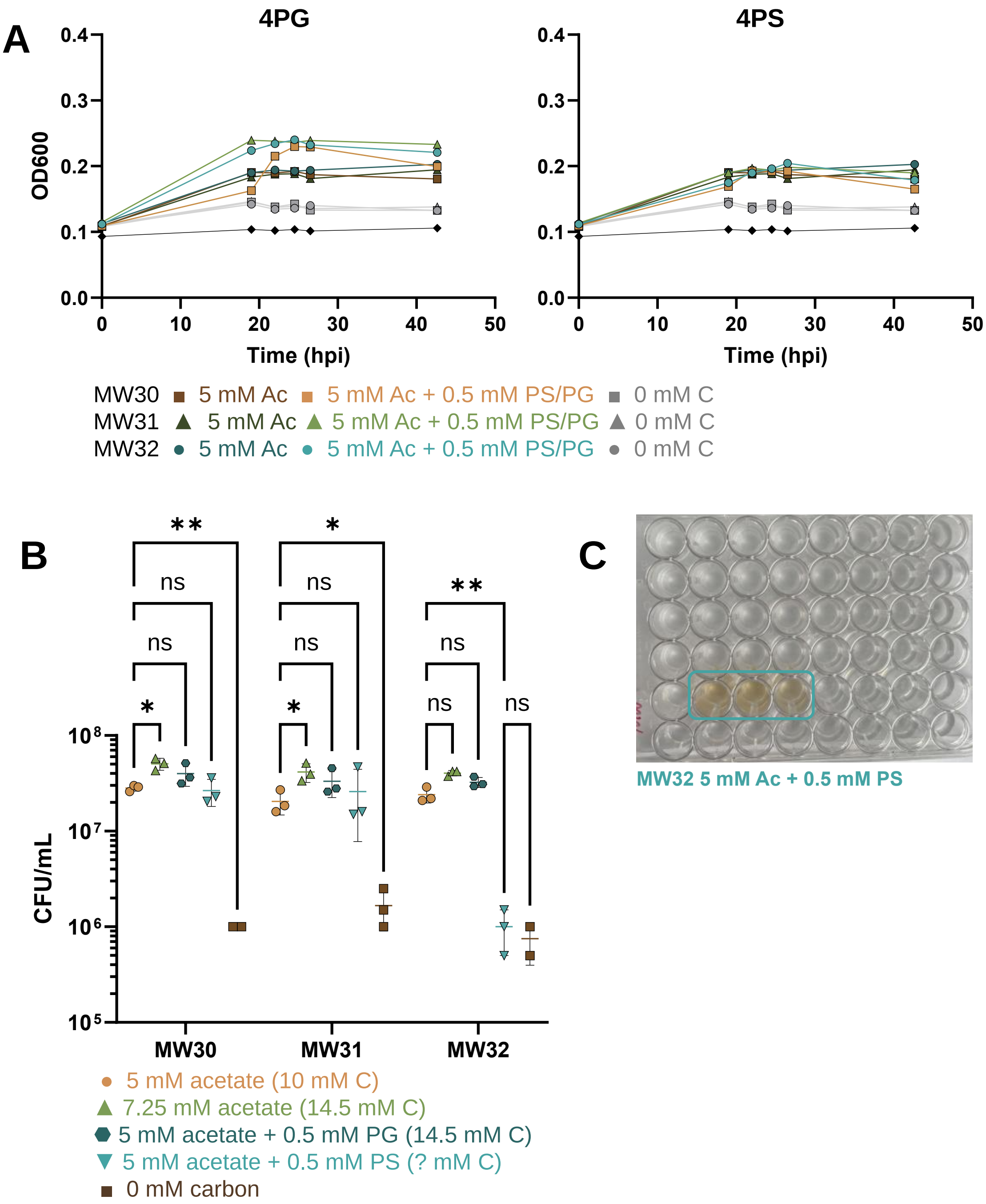


**Figure S11: Growth of Engineered RHA1 on 4PG and 4PS.** OD$_{600}$ of RHAMW30, squares, RHAMW31, triangles, and RHAMW32, circles, grown in media supplemented with 0 mM acetate, grey, 5 mM acetate, darker colors or 5 mM acetate and 0.5 mM of either 4PG **(A)** or 4PS **(B)** Strains were grown in 500 mL M9G media supplemented with the appropriate carbon in a 48 well plate**. (C)** Cells grown on 5 mM acetate and 0.5 mM 4PS turned the growth media a brown color by endpoint (t = 42 h), unlike all other conditions graphed in panels **(A)** and **(B). (D)** CFU in endpoint cultures. Significance testing of a two-way ANOVA: *, *p* = 0.01 - 0.05; **, *p* = 0.001 - 0.01.

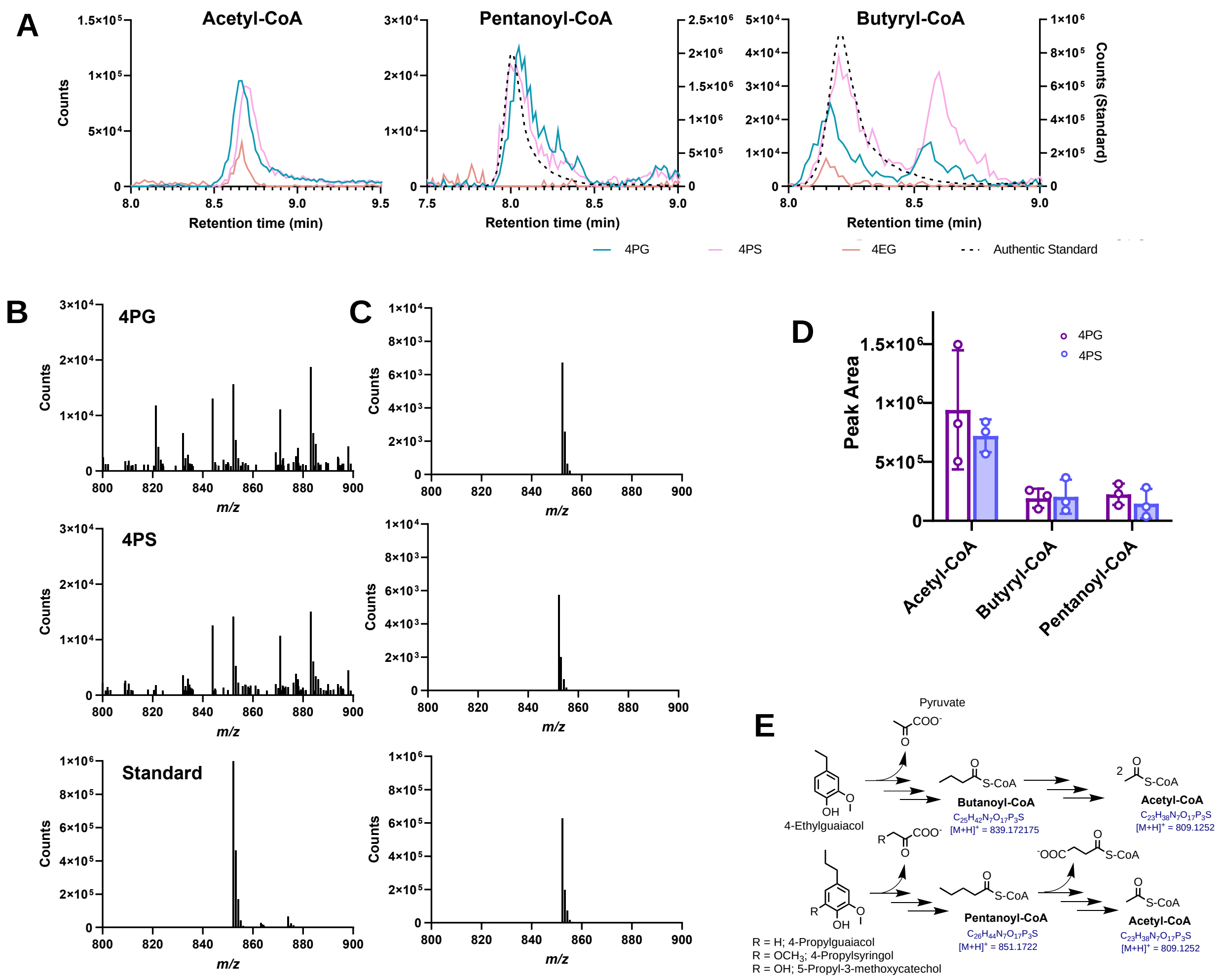


**Figure S12: Acyl-CoAs in cell extracts of RHAMW32 resting cell incubations.** RHAMW32 cells were incubated with 1 mM 4PG or 4PS and ~10 mM glucose. Supernatants were collected after 6 h incubation. (**A**) Predicted pathway of 4EG, 4PG and 4PS to acyl-CoAs, with the predicted protonated *m/z*. (**B**) EICs of predicted *m/z* for acetyl-, pentanoyl- and butyryl-CoA in cell extracts of resting cells incubated with 4 PG (teal), 4PS (orange) and 4EG (pink). Applicable standards (black dotted) are plotted on the right y-axis. 4EG incubated cells were not normalized to the other aromatic substrates (**C**) Peak area of CoAs in cell extracts of resting cells incubated with 4PG (magenta) and 4PS (blue) (**D**) Representative MS spectra of pentanoyl-CoA (LC-MS $t_R$ = 8.1 min) and (**E**) the corresponding subtracted spectra.

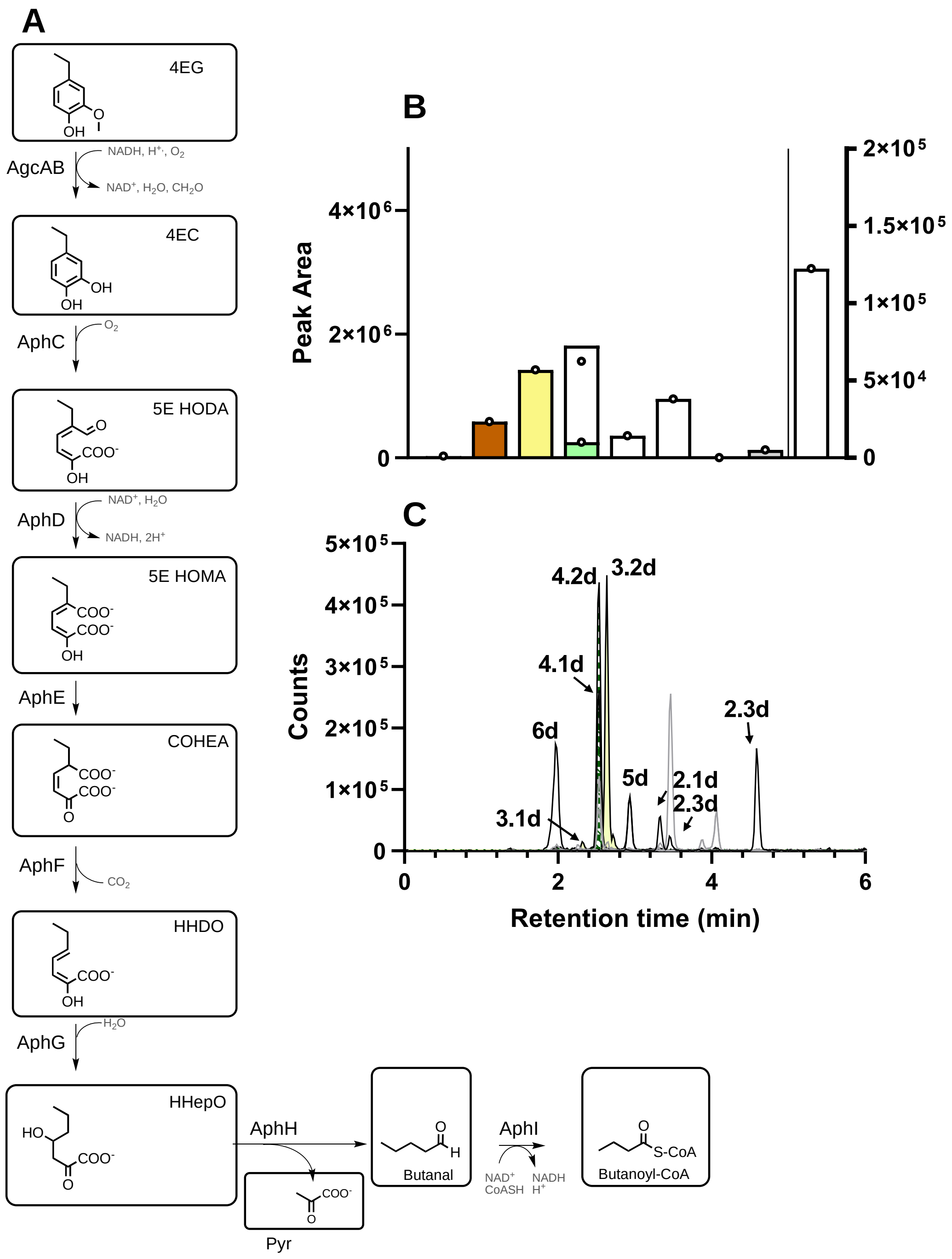


**Figure S13: Catabolism of 4EG by RHA1.** (**A**) Predicted 4EG metabolites. (**B**) EICs of features detected in culture supernatants with *m/z* values of predicted 4PG metabolites (negative mode). RHAMW32 cells were incubated with 0.5 mM 4EG for 2 h. (**C**) Area under the curve of the EICs for the given features. Butanoyl-CoA (But-CoA) was detected in cell extracts. Dashed bars and solid bards indicate the area under the EIC corresponding to the predicted m/z of [M-H2O-H]$^-$, and [M-H]$^-$, respectively. The identities of pyruvate and But-CoA were verified using authentic standards. The other numbered features were provisionally assigned to metabolites based on *m/z* values (**Table S6**).

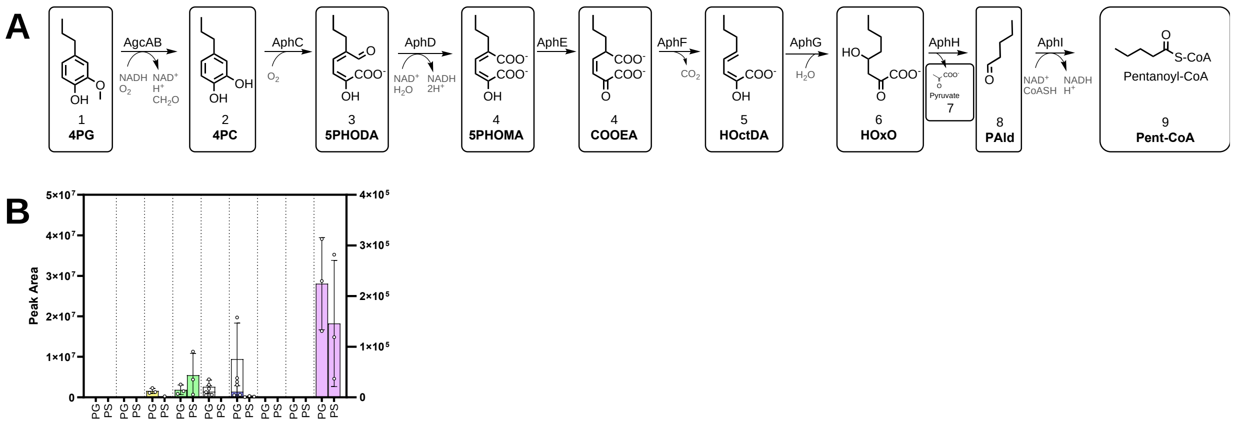


**Figure S14: Supernatant 4PG metabolites of whole cells incubations.** RHAMW32 cells were incubated with 1 mM (magenta bars) 4PG or (blue bars) 4PS and ~ 10 mM glucose. Supernatants were collected after 6 h incubation. (A) Predicted catabolites in 4PG catabolism and (B) Sum of peak area for predicted metabolites in catabolic pathways of 4PG, 4PS, and 4P3MC. Error bars represent the average of three biological replicates.

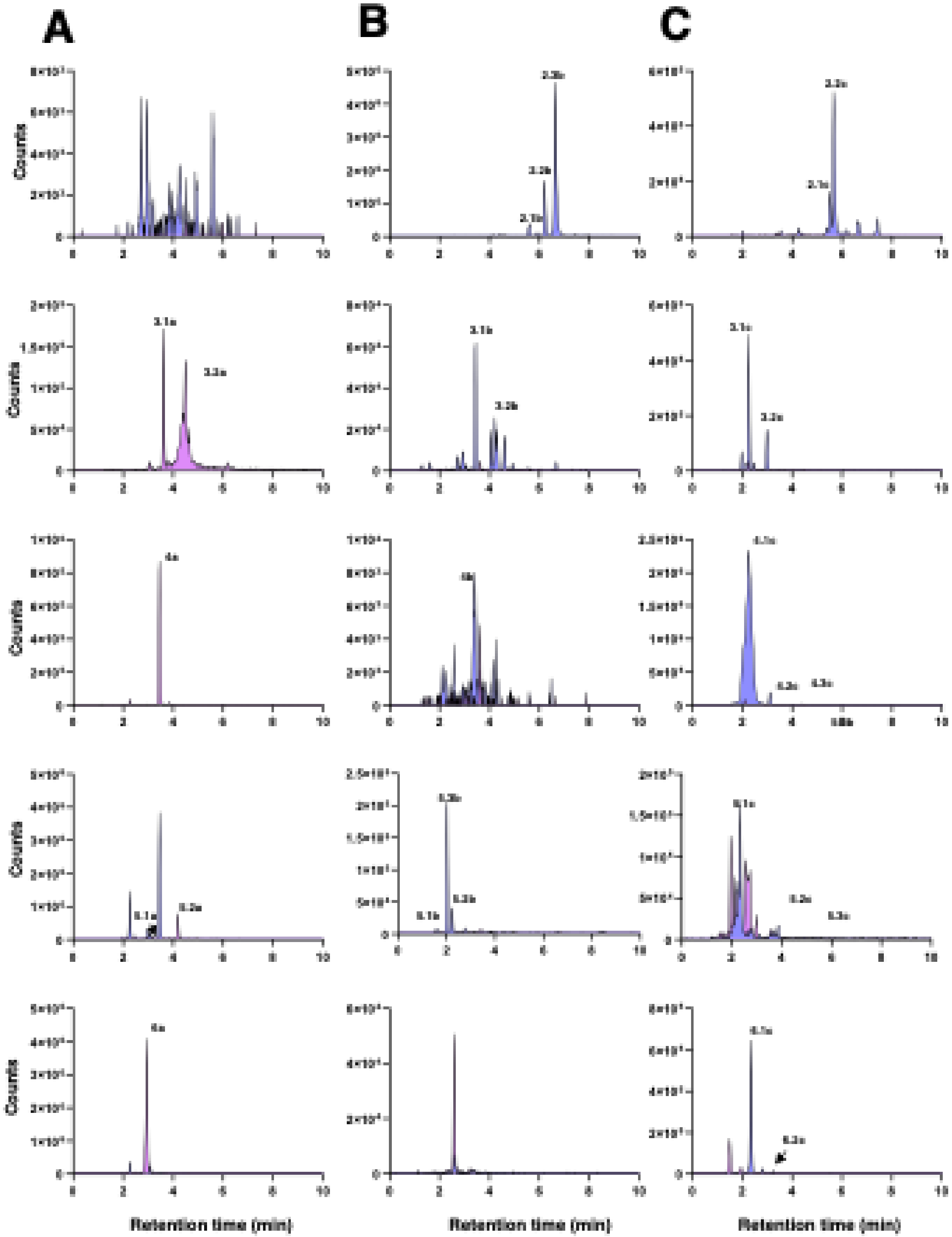


**Figure S15: MS analysis of supernatant metabolites.** Representative (one replicate) EICs of predicted features in the catabolism of (A) 4PG, (B) 4PS and (C) 3M5PC by concentrated RHAMW32 cells. Features **(Table S6)** are indicated above peaks. The EIC of 4PG-incubated cells (magenta) and 4PS-incubated cells (periwinkle) are shown where appropriate.

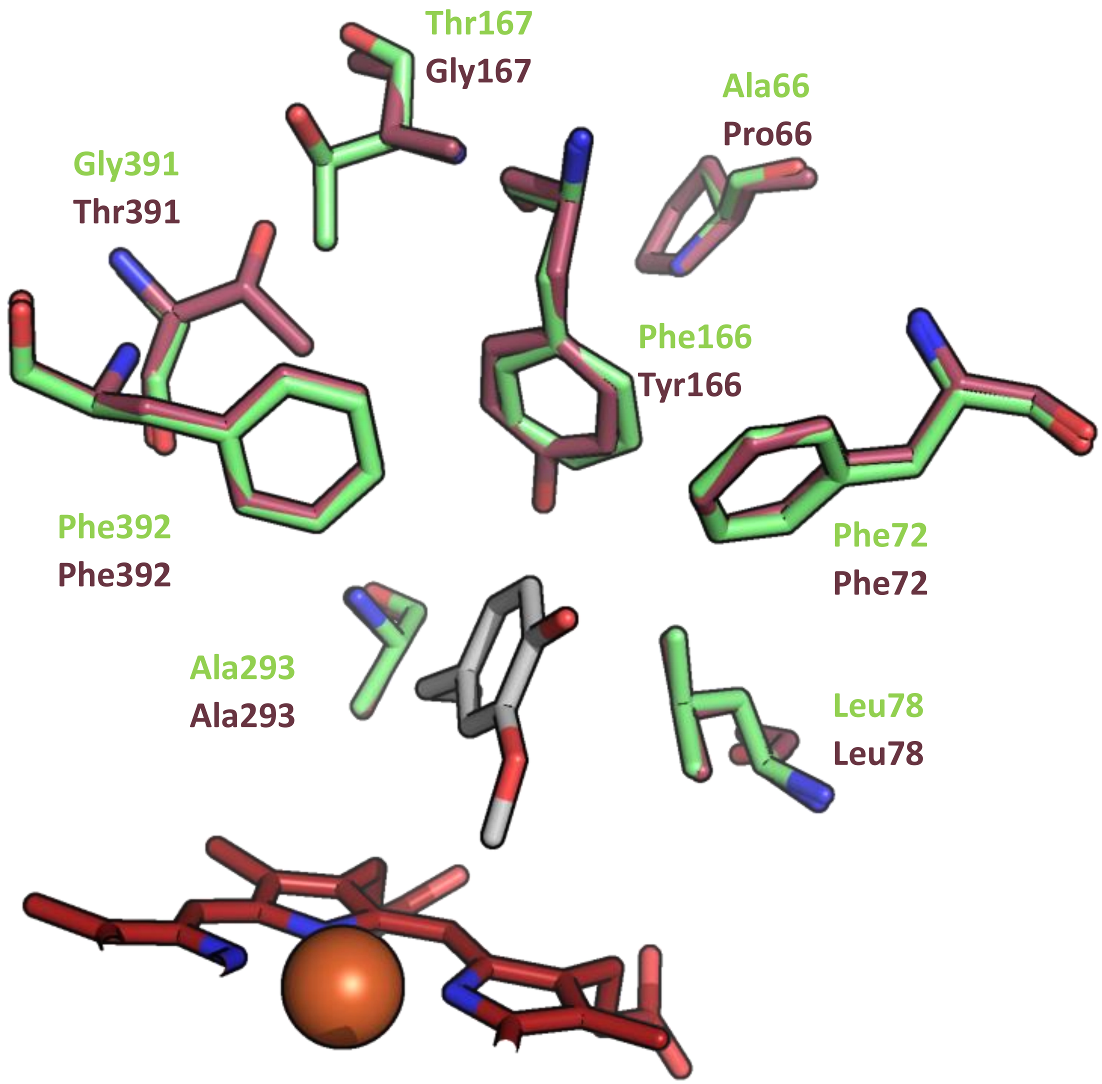


**Figure S16: Comparison of the substrate-binding pockets of the two AgcA homologs**. Overlay AgcA$_{EP4}$ (PDB ID: 9IA1, green) and AgcA$_{RHA1}$ (AlphaFold model, maroon). Side chains of first shell residues are shown along with those of second shell residues 167 and 391. The heme and 4EG carbon atoms are in red and grey, respectively.

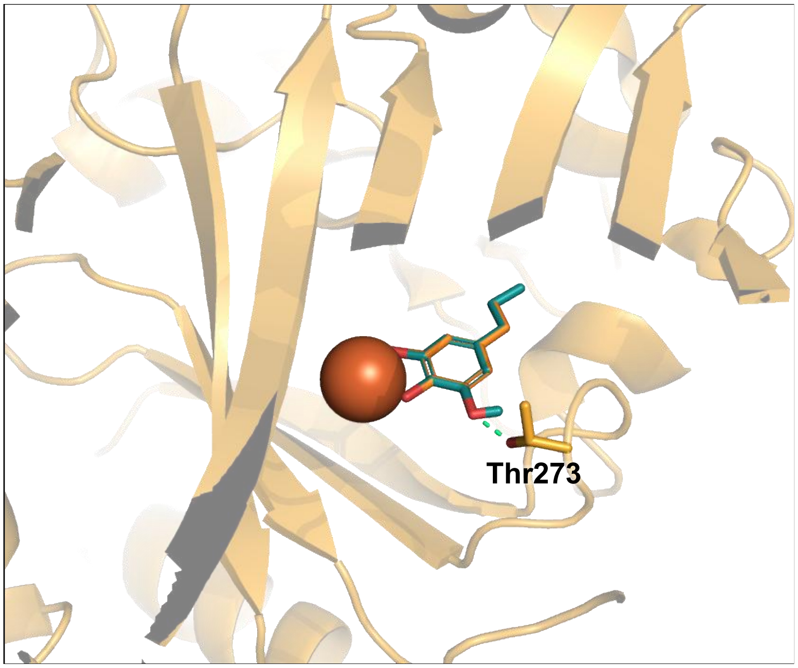


**Figure S17: Binding pocket of AphC$_{RHA1}$ (PBD: 7Q2A) in complex with 4-ethylcatechol.** The predicted *O*-demethylation product of 4PC, 5M3PC (teal) is modelled based on complexed 4EC (orange) position and shown interacting with the active site iron (maroon sphere). Thr273 is highlighted for discussion. The interatomic distance indicated by the aquamarine dashes is 1.9 Å.

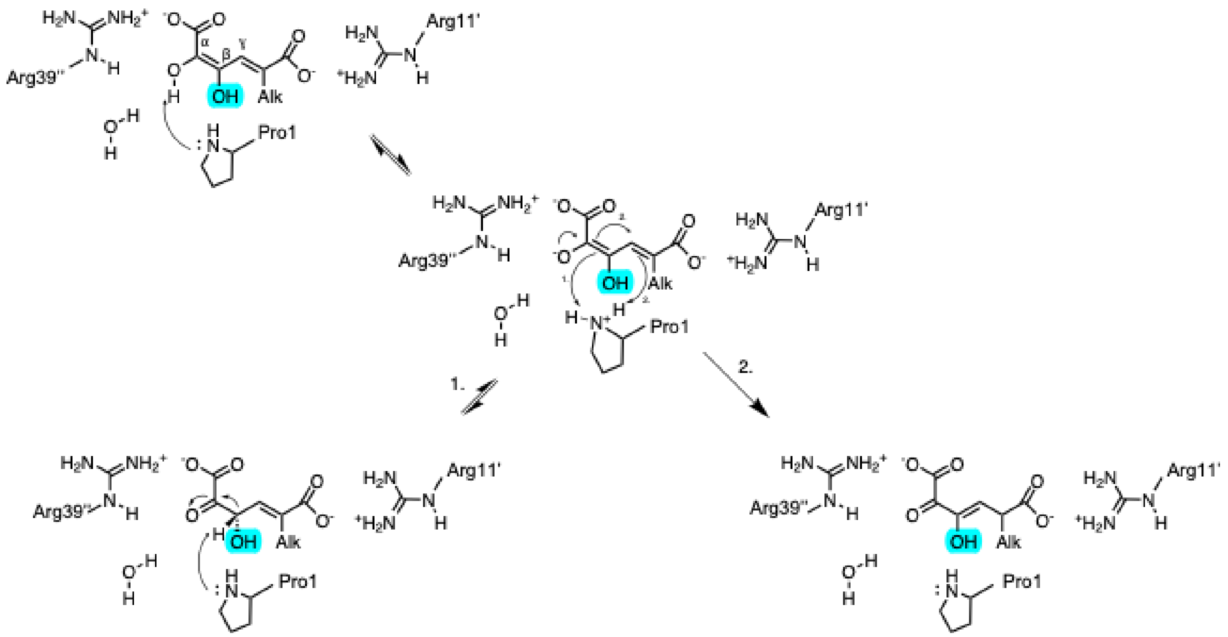


**Figure S18: Proposed mechanism of AphE.** By analogy with homologous 4-oxalocrotonate tautomerases (19), the substrate 3-hydroxy-5-propyl-2-hydroxymuconate (3H5P HOMA) is coordinated by two arginines. Pro1 deprotonates the enol, forming a dienolate which may ttautomerize through one of two paths, forming two keto forms. The substrate of AphE 3-hydroxy-5-carboxy-2-oxooct-3-enoate (3H COOEA) is formed *via* path 2. The additional hydroxyl of the 4PS-derived metabolite is highlighted in turquoise. Adapted from (19).

### TABLES

**Table S1.** Crystallographic diffraction data and refinement statistics of AgcA in complex with 4EG.

| Protein | **AgcA•4EG** |
|---|---|
| PDB code | **9IA1** |
| **Diffraction Data** | |
| Space group | $P\ 4_1\ 2_1\ 2$ |
| Cell dimensions | |
| a, b, c (Å) | 72.56 72.56 348.50 |
| $\alpha$, $\beta$, $\gamma$ (°) | 90 90 90 |
| Resolution [a] | 50.76 - 1.82 (1.89 - 1.82) |
| Observed reflections [a] | 1975487 (186042) |
| Unique reflections [a] | 85027 (8316) |
| Multiplicity [a] | 23.0 (22.4) |
| $I/\sigma(I)$ [a] | 7.63 (0.83) |
| Completeness (%) [a] | 99.91 (99.81) |
| $R_{merge}$ [a] | 0.255 (2.375) |
| $CC_{1/2}$ [a,b] | 0.996 (0.658) |
| **Refinement** | |
| $R_{work}$ (%) [a] | 0.1613 (0.2814) |
| $R_{free}$ (%) [a] | 0.2035 (0.3203) |
| No. of non-hydrogen protein atoms | 7492 |
| No. of ligand atoms | 192 |
| No. of water molecules | 1044 |
| Average B-factor (Å$^2$) | 24.20 |
| RMS Bonds (Å) | 0.013 |
| RMS Angles (°) | 2.068 |
| **Ramachandran** | |
| Preferred (%) | 97.76 |
| Allowed (%) | 2.24 |
| Outliers (%) | 0.00 |
| **Rotamer outliers** | 0.92 |

[a]Values in parentheses are for the highest resolution shell.

[b]Half-dataset correlation coefficient (20).

**Table S2: Binding and activity of $AgcA_{EP4}$ variants on 4PG and 4PS[a]**

| Substrate | Variant | $K_d$[a] | HS[b] | PS depletion[c] | NADH depletion[c] |
|---|---|---|---|---|---|
| | | µM | % | $min^{-1}$ | $min^{-1}$ |
| 4PG | WT | 0.3 (0.1) | >99 | 70 (8) | 67 (6) |
| | F166A | 20 (4) | 20 | 0.324 (0.005) | 2.8 |
| | F166V | 20 (10) | 20 | N.D. | N.D. |
| | F166T | 9 (5) | 10 | N.D. | N.D. |
| | F166N | 12 (2) | 27 | 0.2 (0.1) | 2.8 |
| 4PS | WT | 6.1 (0.9) | 24 | 0.08 (0.04) | 1.8 (0.3) |
| | F166A | 4 (1) | 33 | 0.08 (0.03) | 3.1 |
| | F166V | 3.4 (0.1) | 25 | N.D. | N.D. |
| | F166T | 4.2 (0.6) | 20 | N.D. | N.D. |
| | F166N | 2 (1) | 45 | 0.09 (0.018) | 2.3 |

[a]Error from curve fitting is shown in parentheses.
[b]Percent of heme iron in high spin state in P450:ligand complex.
[c]In the units of µM product $min^{-1}$ µM $heme^{-1}$, simplified to $min^{-1}$. Standard deviation of three replicates is shown in parentheses.

N.D. = Not determined

**Table S3: Coupling of $AgcA_{RHA1}$ WT and Y166A catalyzed *O*-demethylation of 4PS[a]**

| Substrate | Variant | NADH | 4PS | Formaldehyde |
|---|---|---|---|---|
| 4PS | WT | 1.8 (0.3) | 0.8 (0.2) | 0.8 (0.6) |
| | Y166A | 3.9 (0.8) | 3.56 (0.07) | 4.3 (0.2) |

[a]Standard deviation is shown in brackets. All data is in units of µM analyte $min^{-1}$ µM $AgcAB^{-1}$

**Table S4: Rate of substrate turnover of engineered RHA1 strains[a]**

| Strain | 4PG | 4PS |
|---|---|---|
| | **µM $min^{-1}$** | **µM $min^{-1}$** |
| No cell | 0.03 (0.02) | 0.042 (0.008) |
| RHAMW30 | -0.15 (0.02) | 0.00 (0.02) |
| RHAMW31 | -0.147 (0.003) | 0.02 (0.03) |
| RHAMW32 | -0.15 (0.01) | -0.16 (0.02) |

[a]Rates were determined by fitting the linear portion of the depletion curve (t = 30-120 min). Fit errors are in parentheses. Negative rates indicate depletion

**Table S5: *Meta*-cleavage activity of RHAMW32 cell lysates**

| ***Inducer*** | ***Meta-cleavage activity***[a] |
|---|---|
| | **U $g^{-1}$** |
| 4PG | 1.8 (0.5) |
| 4PS | 2.03 (0.02) |
| Glucose | 0.04 (0.05) |

[a]Activity on 4MC normalized to total protein in cell lysates. Standard deviation of three replicates in parentheses.

**Table S6: Metabolites and features in the catabolism of 4PG, 4EG, and 4PS by strain RHAMW32.**

| **Metabolite** | **Abbrev** | **Formula**[a] | **MW**[a] | **Feature** | ***m/z*** | **RT**[b] | **ion** |
|---|---|---|---|---|---|---|---|
| | | | **g/mol** | | | min | |
| 4-propylguaiacol | 4PG | C10H14O2 | 166.22 | **1a** | 165.09 | 5.67 | $[M-H]^-$ |
| 4-propylcatechol | 4PC | C9H12O2 | 152.19 | **2a** | 151.08 | 4.55 | $[M-H]^-$ |
| 2-hydroxy-6-oxo-5-propylhexa-2,4-dienoate | 5P HODA | C9H12O4 | 184.19 | **3.1a**[c] | 183.07 | 3.63 | $[M-H]^-$ |
| | | | | **3.2a**[c] | 183.07 | 4.42 | $[M-H]^-$ |
| 5-propyl-2-hydroxymuconate | 5P HOMA | C9H12O5 | 200.19 | **4.1a**[c] | 199.06 | 2.65 | $[M-H]^-$ |
| | | C9H12O5 | 200.19 | **4.2a**[c] | 181.05 | 3.47 | $[M-H_2O-H]^-$ |
| | | C9H12O5 | 200.19 | **4.3a**[c] | 139.08 | 3.47 | $[M-CO_2 - H_2O- H]^-$ |
| 5-carboxy-2-oxoocta-3-enoate | COOEA | | | | | | |
| 2-hydroxy-octadienote | H OctDA | C8H12O3 | 156.18 | **5a**[c] | 155.07 | 4.2 | $[M--H]^-$ |
| 4-hydroxy-2-oxooctanoate | H OxO | C8H14O4 | 174.20 | **6a**[c] | 173.08 | 2.95 | [M-H]- |
| pyruvate | Pyr | C3H4O3 | 88.06 | **7a** | 87.01 | 1.67 | $[M-H]^-$ |
| pentanal | | C5H10O | 86.13 | | - | - | - |
| pentanoyl-CoA | Pe-CoA | C26H44N7O17P3S | 851.65 | | 852.18 | 8.0 | $[M+H]^+$ |
| 4-ethylguaiacol | 4EG | C9H12O2 | 152.19 | **1d** | | | |
| 4-ethylcatechol | 4EC | C8H10O2 | 138.17 | **2.1d** | 137.06 | 2.65 | $[M-H]^-$ |
| | | | | **2.2d** | 137.06 | 2.80 | $[M-H]^-$ |
| | | | | **2.3d** | 137.06 | 4.58 | $[M-H]^-$ |
| 2-hydroxy-6-oxo-5-ethylhexa-2,4-dienoate | 5E HODA | C8H10O4 | 170.16 | **3.1d**[c] | 169.05 | 2.35 | [M-H]- |

| | | | | | | | |
|---|---|---|---|---|---|---|---|
| 2-hydroxy-6-oxo-5-ethylhexa-2,4-dienoate | 5E HODA | C8H10O4 | | **3.2d**[c] | 169.05 | 2.63 | $[M-H]^-$ |
| 5-ethyl-2-hydroxymuconate | 5E HOMA | C8H10O5 | 186.16 | **4.1d**[c] | 167.03 | 2.53 | $[M-H2O-H]^-$ |
| 5-ethyl-2-hydroxymuconate | 5E HOMA | C8H10O5 | 186.16 | **4.2d**[c] | 167.03 | 2.53 | $[M-CO_2$ $H2O-H]^-$ |
| 5-carboxy-2-oxohept-3-enoate | COHEA | | | | | | |
| 2-hydroxy-heptadienote | HHDO | C7H10O3 | 142.15 | **5d**[c] | 141.06 | 2.93 | $[M-H]^-$ |
| 4-hydroxy-2-oxoheptanoate | HHepO | C7H10O4 | 158.15 | **6d**[c] | 157.05 | 1.98 | [M-H]- |
| pyruvate | Pyr | C3H4O3 | 88.06 | **7a**[c] | 87.01 | 1.67 | [M-H]- |
| Butanal | | C4H8O | 72.11 | | - | - | - |
| Butyryl-CoA | But-CoA | C25H42N7O17P3S | 851.65 | | 852.18 | 8.2 | [M+H]+ |
| 4-propylsyringol | 4PS | C11H16O3 | 196.25 | **1b** | 195.10 | | [M-H]- |
| 5-propyl-3-methoxycatechol | 5P3MC | C10H14O3 | 182.22 | **2.1b** | 181.09 | 5.62 | [M-H]- |
| | | | | **2.2b** | 181.09 | 6.22 | [M-H]- |
| | | | | **2.3b** | 181.09 | 6.63 | [M-H]- |
| 3-methoxy-2-hydroxy-6-oxo-5-propylhexa-2,4-dienoate | 3OMe5P HODA | C10H14O5 | 214.22 | **3.1b**[c] | 213.08 | 3.45 | [M-H]- |
| | | | | **3.2b**[c] | 213.08 | 6.00 | [M-H]- |
| 3-methoxy-5-propyl-2-hydroxymuconate | 3OMe5P HOMA | C10H14O6 | 230.22 | **4b**[c] | 229.07 | 2.73 | [M-H]- |
| 3-methoxy-5-carboxy-2-oxooct-3-enoate | 3OMeCOOEA | | | | 229.07 | 2.73 | [M-H]- |
| 3-methoxy-2-hydroxy-octadienote | 3OMe HOctDA | C9H14O4 | 186.21 | **5.1b**[c] | 185.08 | 2.03 | [M-H]- |
| | | | | **5.2b**[c] | 185.08 | 5.52 | [M-H]- |
| | | | | **5.3b**[c] | 185.08 | 5.67 | [M-H]- |

| | | | | | | | |
|---|---|---|---|---|---|---|---|
| 3-methoxy-4-hydroxy-2-oxooctanoate | 3OMe HOxO | C9H16O5 | 204.22 | **6.1b**[c] | 203.09 | 2.03 | [M-H]- |
| | | | | **6.2b**[c] | 203.09 | 2.25 | [M-H]- |
| 3-methoxy-pyruvate | 3OMePyr | C4H6O4 | 118.09 | | - | - | - |
| Pentanal | | C5H10O | 86.13 | | - | - | - |
| Pentanoyl-CoA | Pe-CoA | C26H44N7O17P3S | 851.65 | | 852.18 | 8.0 | [M+H]+ |
| 5-propylpyrogallol | 5PPG | C9H12O3 | 168.19 | **2.1c**[c] | 167.07 | 5.52 | [M-H]- |
| | | | | **2.2c**[c] | 167.07 | 5.67 | [M-H]- |
| 2,3-dihydroxy-6-oxo-5-propylhexa-2,4-dienoate | 3H5P HODA | C10H14O5 | 214.22 | **3.1c**[c] | 213.08 | 2.26 | [M-H]- |
| | | | | **3.2c**[c] | 213.08 | 3.01 | [M-H]- |
| | | | | **3.3c**[c] | 213.08 | 3.46 | [M-H]- |
| 3-hydroxy-5-propyl-2-hydroxymuconate | 3H5P HOMA | C10H14O6 | 230.22 | **4.1c**[c] | 229.07 | 2.26 | [M-H]- |
| 3-hydroxy-5-carboxy-2-oxooct-3-enoate | 3HCOOEA | | | | | | |
| | 3H5P HOMA | C10H14O6 | 230.22 | **4.2c**[c] | 229.07 | 3.11 | [M-H]- |
| | 3HCOOEA | | | | | | |
| | 3H5P HOMA | C10H14O6 | 230.22 | **4.3c**[c] | 229.07 | 4.43 | [M-H]- |
| | 3HCOOEA | | | | | | |
| 2,3-dihydroxy-octadienote | diHOctDA | C9H14O4 | 186.21 | **5.1c**[c] | 185.08 | 2.35 | [M-H]- |
| | | | | **5.2c**[c] | 185.08 | 3.87 | [M-H]- |
| | | | | **5.3c**[c] | 185.08 | 5.40 | [M-H]- |
| 3,4-dihydroxy-2-oxooctanoate | diHOxO | C9H16O5 | 204.22 | **6.1c**[c] | 203.09 | 2.35 | [M-H]- |
| | | | | **6.2c**[c] | 203.09 | 2.85 | [M-H]- |

| | | | | | | |
|---|---|---|---|---|---|---|
| 3-hydroxy-pyruvate | 3HPyr | C4H6O4 | 118.09 | - | - | - |
| Pentanal | | C5H10O | 86.13 | - | - | - |
| Pentanoyl-CoA | Pe-CoA | C26H44N7O17P3S | 851.65 | 852.18 | 8.0 | [M+H]+ |

[a]Of neutral mass, not necessarily charge state at physiological pH

[b]Retention time on C18 column, except HILIC column for pyruvate (Feature 7a) and CoAs

[c]Provisional assignment, no authentic standard available

**Table S7**: Bacterial strains used in this study

| **Strain** | **Description** | **Source** |
|---|---|---|
| *E. coli* BL-21λ (DE3) | Protein expression | Invitrogen |
| *E. coli* Lemo21 (DE3) | Protein expression | New England Biolabs |
| DH5α | DNA propagation | Invitrogen |
| RHAMW30 | RHA1::pRIME | This study |
| RHAMW31 | RHA1::pRIME-*agcA* | This study |
| RHAMW32 | RHA1::pRIME-*agcA_Y166A* | This study |

**Table S8**: Oligonucleotides used in this study

| Oligo | Description | Sequence (5' to 3') |
|---|---|---|
| pET_agcA_F | Forward Gibson primer for generating pET28-*agcA* | aacctgtattttcagggccatatgaccaccagcaccacg |
| pET_agcA_R | Reverse Gibson primer for generating pET28-*agcA* | ggtggtggtggtgctcgagaagctcagacctcccaggtgac |
| pET_agcB_F | Forward Gibson primer for generating pET28-*agcB* | aacctgtattttcagggccatatgtctgataaatatttggtccgtatagagggaaagaa |
| pET_agcB_R | Reverse Gibson primer for generating pET28-*agcB* | agtggtggtggtggtggtgctcgagtcagagcactgccgccgc |
| pRIME_agcA_F | Forward Gibson primer for generating pRIME-*agcA* | ctttaagaaggagatatacatgtgacacagcaccacgtgg |
| pRIME_agcA_R | Reverse Gibson primer for generating pRIME-*agcA* | cacgggtgccggtgggtcgactagttcagacctcccaggtgac |

**Table S9**: Plasmids used in this study

| Plasmid | Description | Source |
|---|---|---|
| **pET15b** | *E. coli* IPTG inducable expression vector, N-terminal 5His TEV cleavable tag, ampicillin resistance ($\text{Amp}^{R}$), | Novagen |
| **pET15b_*agcA*** | pET15b harboring *agcA* (codon optimized for *E. coli*) | (2) |
| **pET15b_*agcA_A293T*** | pET15b harboring *agcA,* A293T variant | This study |
| **pET15b_*agcA _A293T_L78I*** | pET15b harboring *agcA* A293T_L78I variant | This study |
| **pET15b_*agcA_F166A*** | pET15b harboring *agcA* F166A variant | This study |
| **pET15b_*agcA_F166V*** | pET15b harboring *agcA* F166V variant | This study |
| **pET15b_*agcA_F166T*** | pET15b harboring *agcA* F166T variant | This study |
| **pET15b_*agcA_F166N*** | pET15b harboring *agcA* F166N variant | This study |
| **pET15b_agcA_Y166A** | pET15b harboring *agcA* RHA1 Y166A variant | This study |
| **pET28a** | *E. coli* IPTG inducable expression vector, N-terminal 10-His TEV-cleavable, kanamycin resistance ($\text{Kan}^{R}$) | Modified from Novagen |
| **pET28a_*agcB*** | 10-His pET28a harboring *agcB* EP4 | This study |
| **pET28a_*agcA_RHA1*** | 10-His pET28a harboring *agcA* RHA1 | This study |
| **pET28A_*agcA_RHA1_Y166A*** | 10-His pET28a harboring *agcA* RHA1 Y166A | This study |
| **pRIME_pT1** | Rhodococcal integrative vector, T1 constitutive promoter, ampicillin/apramycin resistance ($\text{Amp}^{R}$/$\text{Apr}^{R}$) | (21) |
